# Phenotypic landscape of a fungal meningitis pathogen reveals its unique biology

**DOI:** 10.1101/2024.10.22.619677

**Authors:** Michael J. Boucher, Sanjita Banerjee, Meenakshi B. Joshi, Angela L. Wei, Manning Y. Huang, Susan Lei, Massimiliano Ciranni, Andrew Condon, Andreas Langen, Thomas D. Goddard, Ippolito Caradonna, Alexi I. Goranov, Christina M. Homer, Yassaman Mortensen, Sarah Petnic, Morgann C. Reilly, Ying Xiong, Katherine J. Susa, Vito Paolo Pastore, Balyn W. Zaro, Hiten D. Madhani

## Abstract

*Cryptococcus neoformans* is the most common cause of fungal meningitis and the top-ranked W.H.O. priority fungal pathogen. Only distantly related to model fungi, *C. neoformans* is also a powerful experimental system for exploring conserved eukaryotic mechanisms lost from specialist model yeast lineages. To decipher its biology globally, we constructed 4328 gene deletions and measured—with exceptional precision--the fitness of each mutant under 141 diverse growth-limiting *in vitro* conditions and during murine infection. We defined functional modules by clustering genes based on their phenotypic signatures. In-depth studies leveraged these data in two ways. First, we defined and investigated new components of key signaling pathways, which revealed animal-like pathways/components not predicted from studies of model yeasts. Second, we identified environmental adaptation mechanisms repurposed to promote mammalian virulence by *C. neoformans*, which lacks a known animal reservoir. Our work provides an unprecedented resource for deciphering a deadly human pathogen.

## INTRODUCTION

This paper concerns a single environmental yeast species, *Cryptococcus neoformans*, which causes ∼20% of deaths in HIV/AIDS^1^. Fungal infections cause 2.5 million annual deaths that disproportionately impact immunocompromised individuals, particularly those with leukocyte defects produced by viral infection, autoantibodies, or the growing number of immunosuppressive medical therapies^2^. Clinically, such infections are among the most difficult to diagnose and treat. Despite the growth of at-risk patient populations and the emergence of new fungal pathogens such as *Candida auris*, antifungal therapy relies on only four drug classes, each with limitations due to toxicity, drug resistance, and/or poor efficacy^3^.

*C. neoformans* is the most common cause of fungal meningitis and sits atop the World Health Organization’s list of priority human fungal pathogens^4^, being responsible for 118,000 deaths each year^1^. Many deaths occur in sub-Saharan Africa, where ∼20% of young adults are HIV-positive in some countries^5^. The case fatality rate is ∼15% in developed countries and much higher in less-resourced settings^6^. *C. neoformans* is found globally in soil, trees, and bird guano^7^, and infection begins by inhalation of spores or desiccated yeast^8^. The resulting pulmonary infection is usually asymptomatic and well-controlled, but in patients with T cell defects, *C. neoformans* disseminates to cause meningitis^9^. Virulence attributes include resistance to stresses (including high temperature), a polysaccharide capsule, phagocytosis suppression, and a secreted factor that enhances type 2 immune responses^8,10^.

A member of the phylum Basidiomycota, *C. neoformans* diverged from ascomycete fungi at least 450 million years ago^11^. Consequently, nearly half of its genes lack orthologs in model yeasts (*Saccharomyces cerevisiae* and *Schizosaccharomyces pombe*), with ∼20% unique to basidiomycetes^12,13^. In addition to its role as a human pathogen, *C. neoformans* is an emerging model organism due to its retention of conserved eukaryotic pathways, including gene silencing and RNA splicing components^14–18^, that have been lost or reduced in model yeasts.

Determining genotype-phenotype relationships for genes under many environmental and/or genetic conditions can help decipher an organism’s biology, in part by placing genes into pathways. Such approaches have been applied at scale to model eukaryotes, such as the ascomycete yeasts *S. cerevisiae* and *S. pombe*, as well as to the protist *Chlamydomonas reinhardtii*, producing global maps and generating hypotheses^19–30^. This approach is more challenging in non-model organisms, including fungal pathogens, as it benefits from genome-scale knockout strain collections and highly precise methods to profile phenotypes. Nonetheless, work on smaller mutant collections in fungal pathogens, including *C. neoformans* and *Candida albicans*, has demonstrated the utility of systematic approaches^31–37^.

Here, we describe the construction and profiling of a genome-scale *C. neoformans* deletion library. Using a high-precision method to measure mutant abundances in pools, we profiled deletion strains in 141 diverse *in vitro* conditions and in a murine infection model, producing a phenotypic landscape of this pathogen. We identified over a hundred functional modules by clustering genes based on their phenotypic signatures. We then leveraged these data to decipher fungal biology of both basic and biomedical importance. First, we identified and investigated new machinery underpinning key pH sensing, stress signaling, and DNA repair pathways. For the latter two, we identified animal-like mechanisms not anticipated from prior studies of model yeasts, demonstrating the utility of global investigations of basidiomycete biology. Second, we identified mechanisms of environmental adaptation that enable mammalian infection, finding prominent roles for glycan biology and secreted enzymes in host colonization and neuroinvasion. Our work illustrates how a high-precision phenotypic landscape can be exploited to decipher pathogen biology at an unprecedented scope.

## RESULTS and DISCUSSION

### Construction of gene deletion collection

We used biolistic transformation^38^ of deletion constructs harboring 1 kb of homology flanking each annotated coding sequencing (CDS) to construct a genome-scale *C. neoformans* knockout library in the haploid KN99α strain background, widely used in the field^39^ (Fig. 1A). We attempted to delete each predicted protein-coding gene at least twice using constructs harboring 1 kb of targeting homology (see Methods). We initially generated 4692 strains. A second round of quality control identified 291 false-positive deletion strains in which the target CDS remained detectable. We also identified 24 strains that had diploidized^40^. Accounting for 49 deletions corresponding to genes no longer present in the current *C. neoformans* H99 genome annotation, this yielded 4328 deletion strains that comprise 62% of the 6975 predicted protein-coding genes (Fig. 1A; Table S1). It has been reported that there are 1465 essential genes in *C. neoformans*^41^, meaning 79% coverage of nonessential genes by our collection. We suspect many putative nonessential genes not deleted correspond to those that produce a significant growth defect when mutated and are therefore considerably more challenging to delete (unpublished observations).

**Figure 1.**
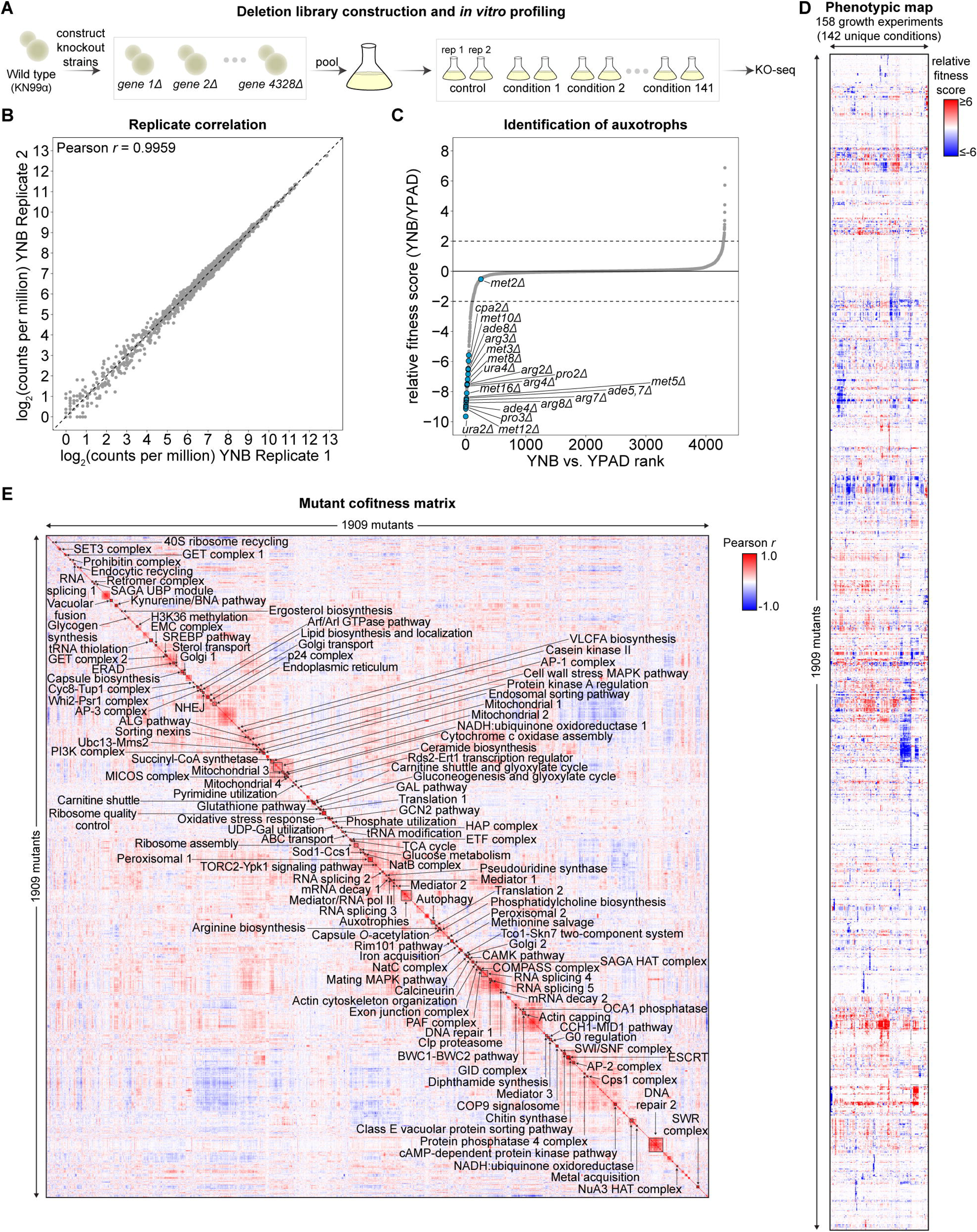
Construction and fitness profiling of a genome-scale deletion library. (A) Overview of deletion library construction and pooled *in vitro* screening. (B) Correlation of KO-seq mutant abundance measurements in replicate YNB cultures. (C) Rank-ordered plot of relative mutant fitness in YNB minimal medium relative to YPAD rich medium. Dashed lines indicate relative fitness scores of −2.0 and 2.0 that were used to define phenotypes. Auxotrophic mutants are highlighted. (D) Phenotypic map. Mutant fitness was measured in 158 growth experiments encompassing 141 unique *in vitro* conditions and 1 *in vivo* condition. Heatmap shows relative fitness scores calculated from endpoint mutant abundances in treated versus same-day control cultures. Data were filtered to show only the 1909 mutants exhibiting at least one condition with relative fitness score ≤ −2.0 or ≥ 2.0 and were clustered based on the centered Pearson correlation (average linkage). (E) Mutant cofitness matrix. Pearson correlation coefficients of phenotypic profiles were calculated and clustered based on the Euclidean distance (centroid linkage). Manually identified clusters are indicated.

### KO-seq enables massively parallel, high-precision fitness measurements

To enable pooled screens of this library, we developed KO-seq, a sequencing-based pipeline to quantify mutant abundances in pools (see Methods). Briefly, knockouts were constructed such that each target gene’s coding sequence was replaced by a nourseothricin acetyltransferase (NAT) drug resistance marker, leaving intact the untranslated regions (UTRs). Following DNA extraction, sequencing-based quantitation of each knockout’s unique NAT-UTR junctions was used to determine the relative abundance of each mutant in a pool. KO-seq measurements of mutant abundances yielded exceptionally precise measurements, with Pearson *r* values typically greater than 0.99 for biological replicates (Fig. 1B). KO-seq identified auxotrophs depleted after growth in minimal (YNB) compared to rich (YPAD) medium (Fig. 1C).

### Fitness profiling of knockout collection

We systematically linked genotype and phenotype by pooling and culturing the entire knockout collection in diverse growth environments and measuring fitness differences with KO-seq (Fig. 1A). We performed 158 pooled fitness experiments, encompassing *in vivo* infection (see below) plus 141 unique *in vitro* conditions corresponding to different media, temperatures, pHs, carbon sources, chemical stressors, and antifungal drugs (Fig. S1A, Table S2). To maximize phenotypes, for each *in vitro* experiment we identified the precise condition that caused detectable growth limitation (see Methods). We identified 1909 mutants with a phenotype—defined as a relative fitness score ≥ 2 standard deviations from the mean—in at least one environment, with a maximum of 99 and a median of 7 phenotypes/mutant (Table S3). Supporting the quality of the data, mutants in known drug targets induced resistance to the corresponding drug, including, for example, flucytosine (*fcy1*Δ, *fcy2*Δ, *fur1*Δ, *uxs1*Δ)^42–44^, FK506 (*frr1*Δ), and rapamycin (*frr1*Δ)^45^, as well as expected genes required for growth in galactose (*gal7*Δ, *gal1*Δ, *uge*2Δ)^46,47^, iron starvation conditions (*hapX*Δ, *hap3*Δ, *hap5*Δ)^48^, and fluconazole (*sre1*Δ, *stp1*Δ, *scp1*Δ)^49,50^ (Fig. S1B-G, Table S3).

### Clustering fitness profiles identifies functional modules

Figure 1D shows all 1909 mutants with at least one phenotype (mean of replicate measurements) as a heatmap after agglomerative hierarchical clustering. Genes that function together in biological pathways share phenotypes, and the spectrum of phenotypes can be used to identify functional genetic modules. A heatmap of the clustered autocorrelation matrix shows correlated mutants clustered along the diagonal (Fig. 1E). We manually annotated clusters in this matrix based on the published literature, orthology to genes of known function in model organisms, and bioinformatic prediction of protein functional domains, identifying 127 gene modules (Fig. 1E). Representative clusters are shown in Figure S2; these include the NAD biosynthesis (kynurenine) pathway^51^ (Fig. S2A); the SREBP hypoxia adaptation pathway^49,50,52–54^ (Fig. S2B); the protein kinase A pathway ^55–60^ (Fig. S2C); and polysaccharide capsule genes^61–63^ (Fig. S2D). Additional clusters, including potentially novel modules, are shown in Fig. S2E-Z, S2AA. A full set of annotated modules can be found in Table S4. Below, we describe how investigations of these modules can yield biological insights.

### Insights into a Hedgehog-like pH-sensing pathway

Fungi harbor a conserved pH sensing pathway, the Rim 101 pathway, that senses high pH and is required for adaptation to this environment^64^ (Fig. 2A). The precise mechanisms of pH sensing are unknown. In *C. neoformans,* pH has been proposed to be sensed by seven-transmembrane protein Rra1^65^ (<u>R</u>equired for <u>R</u>im101 <u>a</u>ctivation 1); its activation results in the high pH-dependent assembly of surface puncta by the downstream signaling component Rim23^66^. Rim23 is part of a protease complex containing the adaptor Rim20 and the calpain proteinase Rim13 that cleaves and activates Rim101, a GLI family transcription factor^67–69^. Cleavage enables nuclear entry of Rim101 and activation of pH-responsive genes, enabling survival at high pH^67–69^. As seen for the activation of the mammalian Rim13 ortholog calpain-7 *in vitro*^70–72^, ESCRT proteins play a noncanonical role in activating the Rim13 calpain *in vivo*^67,68^. Because animal GLI family protein transcription factors are also activated for nuclear entry by proteolytic cleavage in response to cell surface signals during Hedgehog signaling^73^, similarities between the two pathways have been noted^74^, although the mechanism of proteolysis appears to have been rewired^75^.

**Figure 2.**
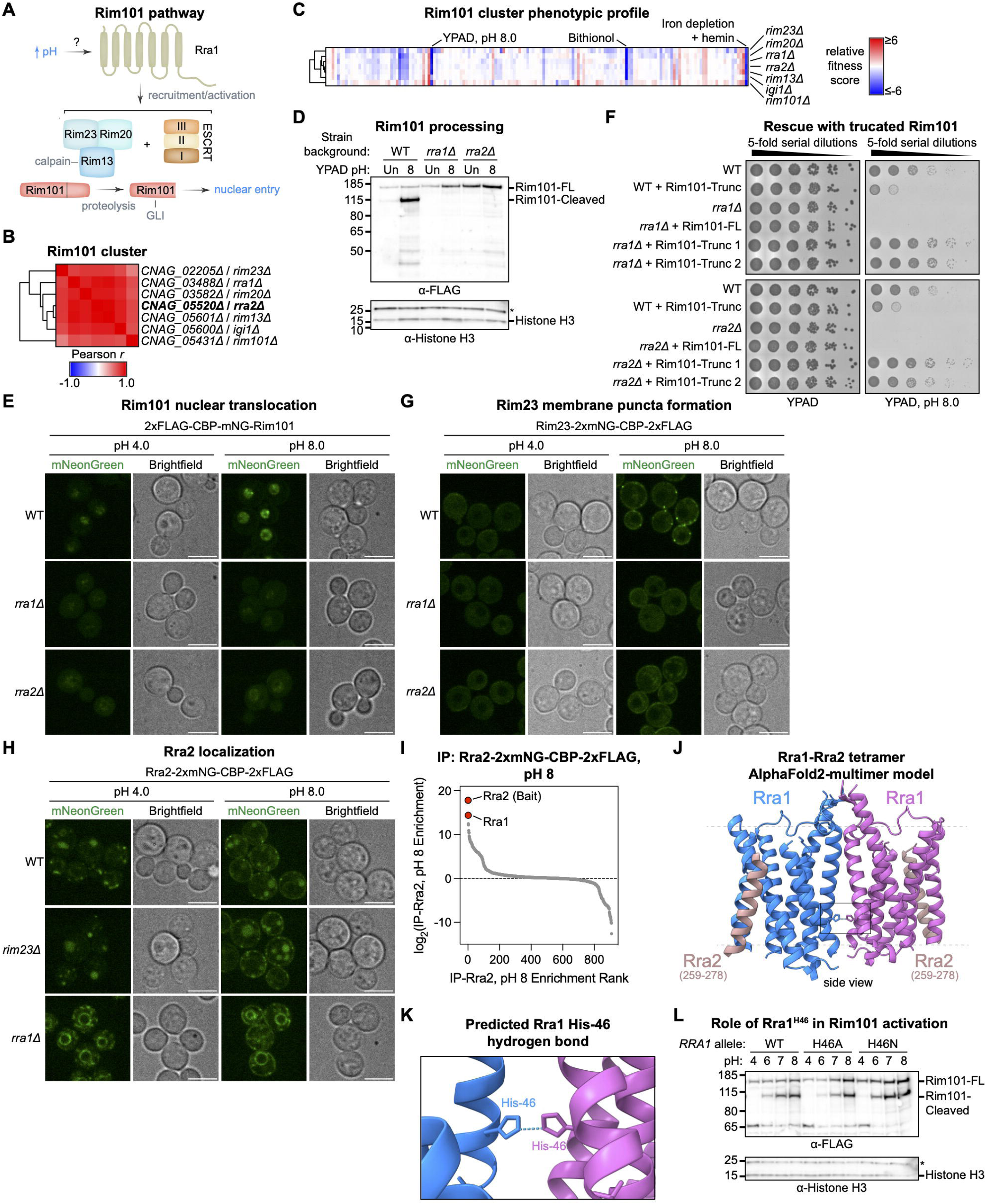
Insights into a Hedgehog-like pH sensing pathway. (A) Diagram of the Rim101 pH sensing pathway in *C. neoformans*. (B) Rim101 pathway cluster. (C) Phenotypic profile of Rim101 pathway members. (D) Western blot assaying proteolytic processing of endogenously tagged 2xFLAG-mNG-CBP-Rim101 in unbuffered medium or at pH 8.0. Asterisk indicates nonspecific band. (E) Live confocal imaging of endogenously tagged 2xFLAG-CBP-mNG-Rim101 in SC medium buffered to pH 4.0 or 8.0 with McIlvaine’s buffer. (F) Spot dilution assay assessing rescue of Rim101 pathway mutants by expression of a truncated Rim101 allele. Images were taken after 3 days of growth. (G) Live confocal imaging of endogenously tagged Rim23-2xmNG-CBP-2xFLAG in SC medium buffered to pH 4.0 or 8.0 with McIlvaine’s buffer. (H) Live confocal imaging of endogenously tagged Rra2-2xmNG-CBP-2xFLAG in SC medium buffered to pH 4.0 or 8.0 with McIlvaine’s buffer. (I) Rra2 AP-MS at pH 8. YPAD cultures of yeast expressing endogenously tagged Rra2-2xmNG-CBP-2xFLAG were shifted to pH 8 for 1 hour, and anti-FLAG AP-MS was performed on membrane extracts. Data represent the protein-level ratio of normalized MS1 area from tagged versus untagged strains grown and processed in parallel. Averaged from 2 biological replicates. See also Figure S3E. (J) AlphaFold2-multimer model of predicted Rra1-Rra2 tetramer. See also Figure S3F-H. Confidently predicted protein segments are displayed. (K) Inset from (J) showing a predicted Rra1^H^^46^-Rra1^H^^46^ hydrogen bond. (L) Western blot assaying proteolytic processing of endogenously tagged 2xFLAG-mNG-CBP-Rim101 in response to *RRA1* alleles encoding His46 mutants. Asterisk indicates nonspecific band. See also Figure S3I.

A module harboring components of the Rim101 pathway contains a previously unannotated gene, which we name *RRA2*, encoding a protein with two predicted transmembrane domains (Fig. 2B, C – *IGI1* in the cluster is adjacent to *RIM13* and was not considered further). We confirmed that *rra2*Δ cells fail to grow at high pH and that this phenotype could be complemented by the expression of the wild-type protein (Fig. S3A, B). Cells lacking Rra2 are defective in proteolytically processing Rim101 upon a shift to alkaline pH (Fig. 2D), and Rim101 does not accumulate in the nucleus under these conditions, in contrast to wild-type strains (Fig. 2E). As with other *bona fide* pathway components, the expression of a C-terminally truncated *RIM101* allele rescues the alkaline pH sensitivity of strains lacking Rra2 (Fig. 2F).

Like Rra1, Rra2 is necessary for recruitment of Rim23 to membrane puncta following shift to alkaline pH (Fig. 2G), suggesting that it acts upstream of the Rim20-Rim23-Rim13 complex. Because Rra1 and Rra2 both contain predicted membrane-spanning domains and act as upstream components of the pathway, we investigated their relationship. Rra2 is not required for Rra1 accumulation in surface puncta following a pH shift (Fig. S3C), but Rra1 is required for Rra2 accumulation in surface puncta; in the absence of Rra1, Rra2 displays a localization pattern characteristic of ER proteins^76^, indicating a trafficking defect (Fig. 2H). Rim23 is not required for the localization of Rra2 (Fig 2H). The simplest interpretation of these findings is that Rra1 and Rra2 act together at the membrane to initiate pH signaling.

To investigate the pathway further, we applied AlphaFold2-multimer^77^ to all pairwise combinations of Rim pathway components. As shown in Fig. S3D, we obtained predictions for pairwise interactions between multiple components of the pathway. In the case of Rra2, AlphaFold2-multimer predicts interactions with Rra1 and Rim23. Using a FLAG-tagged allele of Rra2, we performed affinity purification-mass spectrometry (AP-MS) at both low and high pH. Because Rra2 is a predicted membrane protein, we extracted a membrane pellet with detergent (see Methods) prior to purification with anti-FLAG antibodies and mass spectrometry. Rra1 was the most highly enriched protein (as defined by MS1 area signal) relative to control extracts from an untagged strain (Fig. 2I, S3E, Table S5), providing biochemical evidence that Rra2 physically associates with its functional partner Rra1.

We noticed a high-confidence prediction for an Rra1 dimer, which was not mutually exclusive with the predicted Rra1-Rra2 complex (Fig. 2J, S3F-H). Analysis of the Rra1-Rra1 dimer interface revealed geometry feasible for hydrogen bonding between His46 residues on each monomer (Fig. 2J, K). Because histidine is the only amino acid with a side-chain pK_a_ near neutral^78,79^, and pH-sensitive histidine-histidine hydrogen bonding between two mono-protonated (as opposed to di-protonated) side-chains is known to occur^80^, we tested whether His46 played a role in pH sensing. We generated strains harboring Rra1 alleles with mutations expected to disrupt (H46A) or stabilize (H46N) interchain chain hydrogen bonding, the latter predicted to introduce pH-independent hydrogen bonds between the amide nitrogen and oxygen moieties of corresponding Asn46 residues. Strikingly, strains expressing Rra1^H46A^ displayed reduced pH-dependent signaling, as measured by Rim101 processing, whereas strains expressing Rra1^H46N^ displayed increased signaling (Fig. 2L, Fig. S3I). Both mutations had partial effects, consistent with recent saturation mutagenesis of a mammalian proton-sensing GPCR showing that pH-sensing activity is distributed among multiple residues, including several histidines^81^. The data are consistent with the hypothesis that high pH promotes His46-dependent Rra1 dimerization to activate the pathway; however, definitive tests of this model will require structural work. Nonetheless, this finding begins to crack open the biochemical mechanism of pH-controlled cleavage of a GLI family transcription factor.

### Fungal orthologs of animal UNC79 and UNC80 activate calcineurin signaling

In animals, neuronal excitability is determined by the resting membrane potential (RMS). RMS is controlled by channels, particularly the voltage-insensitive NALCN sodium leak channel complex, which contains four major subunits: the membrane protein NALCN α1 subunit and the accessory proteins FAM155A, UNC79, and UNC80^82^. UNC79 and UNC80 are giant HEAT-repeat proteins that form a heterodimer associated with the cytoplasmic face of the NALCN complex^82^. Mutations in NALCN, UNC79, and UNC80 underpin several inherited diseases and have been proposed to be involved in many others^82^. Surprisingly, NALCN and FAM155A are ancestrally related to the fungal proteins Cch1 and Mid1, respectively, indicating that the proteins evolved before animals and neurotransmission^83,84^. Cch1 and Mid1 form a calcium channel that acts in a stress survival pathway^85–89^ upstream of the calcium-activated phosphatase calcineurin, the target of the clinically used immunosuppressants cyclosporine and FK506, which arrest fungal growth^90,91^ (Fig. 3A). The fungal transcription factor Crz1, first identified in *S. cerevisiae* but also present in *C. neoformans*, is analogous to the mammalian NFAT protein (critical for T cell activation^92–94^) and, like NFAT after T cell receptor ligation^95^, is dephosphorylated by calcineurin, enabling its nuclear entry^96–102^ (Fig. 3A). This linear model is an oversimplification because *C. neoformans cch1*Δ and *mid1*Δ knockout strains are viable at 37°C^103^ while calcineurin knockouts fail to grow under this stress condition^104^, suggesting Cch1-Mid1-independent sources of Ca^++^. Furthermore, in *S. cerevisiae,* there is evidence for calcineurin-dependent feedback inhibition of Cch1-Mid1^85,105^, which presumably mitigates the deleterious effects of excessive cytosolic Ca^++^ accumulation (Fig. 3A).

**Figure 3.**
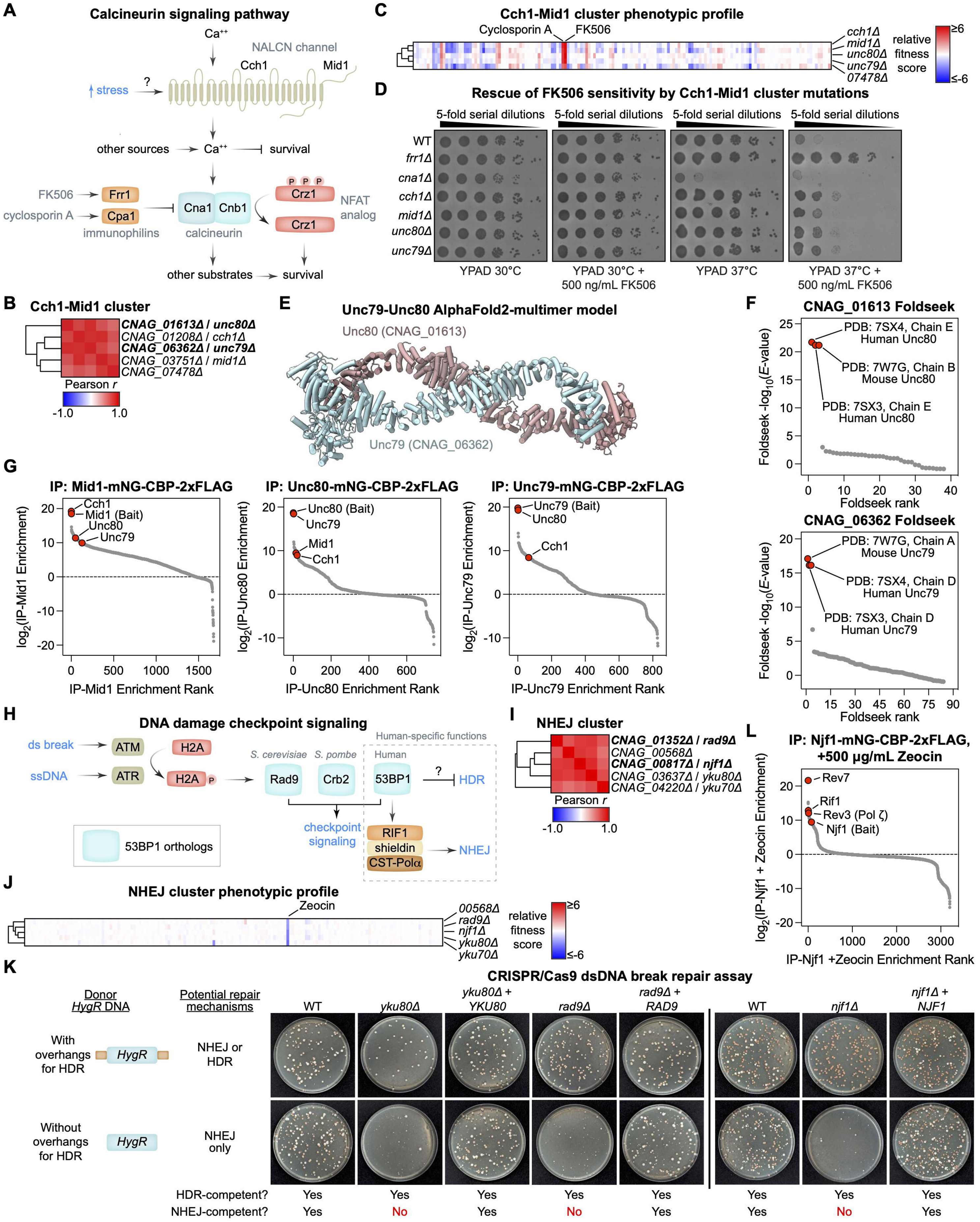
Mammalian-like signaling pathways in *C. neoformans*. (A) Diagram of calcineurin signaling pathway in *C. neoformans*. (B) Cch1-Mid1 cluster. (C) Phenotypic profile of Cch1-Mid1 cluster members. (D) Spot dilution assay assessing sensitivity of Cch1-Mid1 cluster mutants to FK506 at 30°C and 37°C. Images were taken after 2 days of growth. See also Figure S4A. (E) AlphaFold2-multimer model of predicted CNAG_06362-CNAG_01613 (Unc79-Unc80) dimer. See also Figure S4B. Confidently predicted protein segments are displayed. (F) Foldseek results from PDB100 database search with CNAG_01613 and CNAG_06362 AlphaFold2 models from (E). (G) Mid1, Unc79, and Unc80 AP-MS. Anti-FLAG AP-MS was performed on membrane extracts from strains expressing endogenous *C*-terminal mNG-CBP-2xFLAG tags on the indicated strains. Data represent the protein-level ratio of normalized MS1 area from tagged versus untagged strains grown and processed in parallel. Averaged from 2 biological replicates. (H) Diagram of DNA damage checkpoint signaling. (I) NHEJ cluster. (J) Phenotypic profile of NHEJ cluster members. (K) Rescue of Cas9-induced double-stranded breaks by a drug-resistance donor template with or without overhangs for HDR. The indicated strains were transformed with PCR products to express 1) Cas9; 2) an sgRNA targeting the *ADE2* gene; and 3) a hygromycin B selection marker (*HygR*) with or without overhangs for HDR. See Figure S5B for diagram. Transformants were selected on YPD plates containing hygromycin B. Strains deficient in NHEJ are expected to repair the Cas9-induced double-stranded break when overhangs for HDR are supplied (top row) but not when they are omitted (bottom row). Pink/red colony color occurs due to accumulation of purine precursors in the absence of a functional *ADE2* gene. (L) Njf1 AP-MS following Zeocin treatment. YPAD cultures of yeast expressing endogenously tagged Njf1-mNG-CBP-2xFLAG were treated with 500 μg/mL Zeocin for 2 hours, and anti-FLAG AP-MS was performed on clarified cell lysates. Data represent the protein-level ratio of normalized MS1 area from tagged versus untagged strains grown and processed in parallel. From 1 biological replicate. See also Figure S5C.

A cluster containing knockouts corresponding to *CCH1* and *MID1* harbored knockouts in two additional genes, *CNAG_01613* and *CNAG_06362*, which encode large (>2000 amino acids) previously unstudied proteins (Fig. 3B; *CNAG_07478* in the cluster lies adjacent to *MID1* and is not considered further). The primary phenotype of mutants in this cluster is quantitative tolerance to FK506 and cyclosporin A (Fig. 3C). The phenotype was considerably more dramatic at 37°C, a temperature at which calcineurin becomes essential and FK506 and cyclosporin A inhibit growth completely (Fig. 3D). These phenotypes were recapitulated in independently constructed knockouts and could be complemented by the wild-type genes (Fig. S4A).

While CNAG_01613 is unannotated, algorithms suggested it contains HEAT repeats (also known as Armadillo repeats) and an UNC80 C-terminal motif hit. CNAG_06362 is predicted to contain a protein kinase C-like diacylglycerol binding domain. We generated AlphaFold2-multimer models of the two proteins and observed the formation of helical HEAT repeat monomers that form an antiparallel heterodimer (Fig. 3E; Fig. S4B). When we searched for structural homologs of the AF-multimer prediction using Foldseek^106^, by far the most significant hits were to the UNC79 and UNC80 structures observed in the cryoEM structures of the mammalian NALCN channelosome (Fig. 3F). We renamed the genes accordingly.

To test for biochemical interactions, we endogenously tagged Mid1, Unc79, and Unc80 with a 2XFLAG epitope and performed AP-MS from solubilized membrane pellets. The most enriched protein in the Unc79 sample was Unc80 and vice-versa (Fig. 3G, Table S5), consistent with the prediction of a heterodimer. As expected, the top hit for the Mid1 purification was Cch1 (Fig. 3G, Table S5). Likewise, Unc79 and Unc80 were enriched in the Mid1 purification, both Mid1 and Cch1 were enriched in the Unc80 purification, and Cch1 was enriched in the Unc79 purification (Fig. 3G, Table S5).

Unc79 and Unc80 were previously thought to be animal-specific^84^. However, our genetic data, AlphaFold2 predictions, and biochemical data support the existence of a mammalian-like NALCN channelosome complex in *C. neoformans*, indicating that the entire complex was present in the common ancestor of fungi and animals. We did not detect orthologs of the *C. neoformans* Unc79/Unc80 proteins outside of basidiomycetes, suggesting that either the proteins were lost or highly diverged in ascomycetes such as *S. cerevisiae*. Thus, *C. neoformans* provides a system for dissecting the precise molecular function of UNC79 and UNC80 in channel function, which remains poorly understood. Finally, our findings of resistance to calcineurin-targeting drugs via mutations in Unc79/Unc80 are germane to efforts to develop fungal-specific inhibitors of calcineurin^107^.

### A fungal 53BP1/Shieldin-like pathway promotes nonhomologous end-joining

53BP1, an animal protein identified as a binding partner for the p53 tumor suppressor, promotes non-homologous end joining (NHEJ) of double-strand breaks over homology-directed repair (HDR)^108,109^. To promote NHEJ, 53BP1, together with RIF1, recruits the Shieldin complex, which consists of SHLD1, SHLD2, SHLD3, and REV7^110–113^. Shieldin, in turn, recruits the fill-in DNA polymerase CST-Polα-primase^112^ to counter single-stranded DNA resection, which would inhibit the ligation of double-strand breaks. Pol ζ has recently been shown to function in the Shieldin pathway, presumably to further fill-in synthesis^114^ (Fig. 3H).

Fungal orthologs of 53BP1 include Rad9 in *S. cerevisiae* and Crb2 in *S. pombe*^115–117^. Both contain Tudor and C-terminal BRCT domains, and the *S. pombe* ortholog’s Tudor domain, like that of 53BP1, recognizes H4-K20me2^118^. However, unlike 53BP1, the *S. cerevisiae* and *S. pombe* 53BP1 orthologs are not known to be part of a RIF1-Shieldin-CST-Polymerase α/ζ pathway. Moreover, aside from Rev7, which is part of multiple complexes, Shieldin subunits are not detectable outside of vertebrates (unpublished observations).

Thus, we were surprised to observe that a knockout of 53BP1/Rad9/Crb2 ortholog in *C. neoformans*, which we named Rad9, and a novel protein encoded by *CNAG_00817*, cluster with knockouts of critical NHEJ factors, the Ku complex proteins (Fig. 3I). Knockouts in this cluster display sensitivity to the double-strand break-inducing compound Zeocin, indicating a crucial role for NHEJ in surviving DNA breaks in *C. neoformans* (Fig. 3J). Independently generated knockouts of *RAD9*, *CNAG_00817,* and the two Ku orthologs, *YKU70* and *YKU80*^119^, recapitulate the Zeocin sensitivity of the original mutants (Fig. S5A). The uncharacterized gene *CNAG_00568* in this cluster did not validate (Fig. S5A).

Next, we tested whether *C. neoformans RAD9* and *CNAG_00817* were required for NHEJ. We recently observed that NHEJ can be assayed by transforming a Cas9-expressing strain^76^ with DNA molecules encoding an sgRNA and a selectable marker lacking genome homology. Selected colonies near-invariably carry an insertion of the marker DNA at the break site by ligation (M. Huang et al., in preparation). Mutants lacking the Ku proteins display a near-complete defect in the transformation frequency of such a construct (M. Huang et al., in preparation). Strains deleted for *C. neoformans RAD9* or *CNAG_00817* also display a strong defect in this NHEJ-dependent repair assay (Fig. 3K, S5B). In contrast, donors containing homology arms produced transformants, indicating that HDR is intact in these mutants (Fig. 3K). We named *CNAG_00817* to *NJF1* for *N*onhomologous end *j*oining *f*actor *1*.

To gain additional insight, we performed AP-MS experiments on strains harboring FLAG-tagged Rad9 or Njf1, which were treated or not with Zeocin. Strikingly, Njf1 copurified with the *C. neoformans* orthologs of Rev7, Rif1, and Rev3/Pol ζ (Fig. 3L, S5C, Table S5). No clear interactors were identified for Rad9 (Table S5). Since Rev7 is a *bona fide* component of the Shieldin complex and Rif1 an adaptor that recruits it, Njf1 displays functional properties and biochemical associations analogous to those of Shieldin. Consistent with this model, AlphaFold2-multimer produced a high-confidence Njf1-Rev7 complex prediction (Fig. S5D). Moreover, the prediction shows a peptide from Njf1 captured by the HORMA domain of Rev7 and its ‘seatbelt’ in a manner typical of other Rev7 ligands, including SHLD3^120,121^. Deepening the analogy with Shieldin is the recent discovery that Rev3/ Pol ζ, which we find associated with Njf1, is an effector of Shieldin^114^. We conclude that 1) the 53BP1-dependent promotion of NHEJ is far more ancient than previously supposed but lost in *S. cerevisiae* and *S. pombe* and 2) a novel Shieldin-like pathway requiring Njf1 exists in *C. neoformans*. Because Njf1 orthologs are only detectable only in basidiomycetes (unpublished observations), tracing the origins and ancestry of this pathway outside of this phylum will require biochemical and functional experiments.

### Insights into pathogen mechanisms that enable fitness in the host

To identify genes required for infectivity, we generated pools of up to 480 knockouts and inoculated each pool into five replicate C57BL/6J mice intranasally (Fig. 4A). Upon reaching their endpoint, animals were sacrificed, and relative mutant abundances in the lung were quantified by KO-seq (see Methods). Mutant abundances in replicate mice were reproducible, with median Pearson *r* = 0.90 (Fig. S6A and S6B). We calculated fitness scores derived from the standardized log_2_ fold changes in mutant abundance (see Methods). We classified 574 mutants with an average fitness score of less than −2.0 as having *in vivo* fitness defects, as well as 36 mutants that were modestly enriched in infected mice (Fig. 4B; Table S6). These data significantly overlapped with prior data using a smaller mutant library in a different *C. neoformans* strain analyzed in a different mouse strain background^32^ (Fig. 4C). We re-pooled and retested the mutant in mice and observed that >98% re-tested (Fig. 4D). While many genes required for infection are orthologous to components of critical cellular pathways in model systems, 84 genes (15%) are unique to basidiomycetes (Fig. 4E, S6C-E).

**Figure 4.**
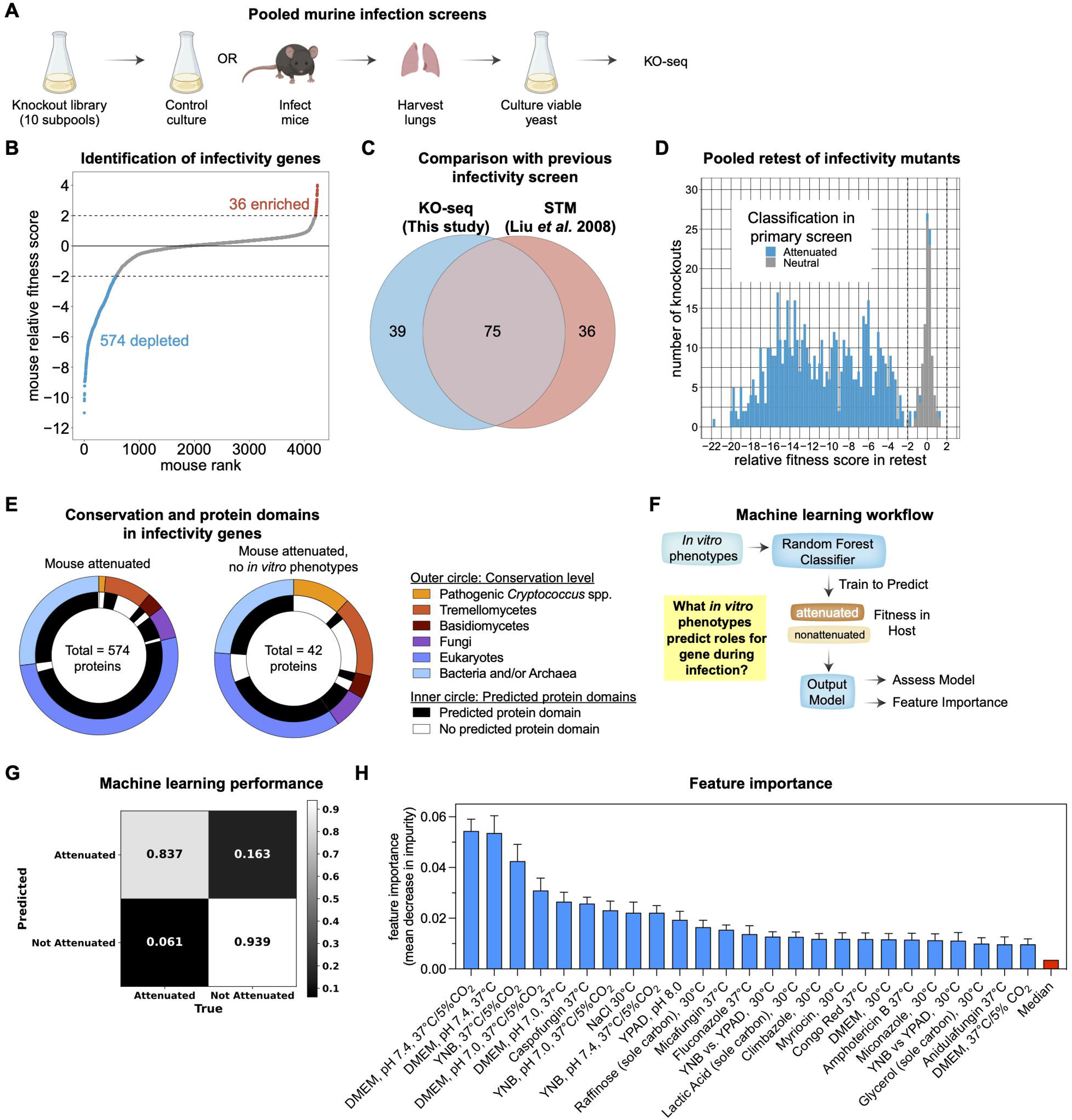
Pathogen functions that promote mammalian infection. (A) Schematic of *in vivo* fitness screens in a murine pulmonary infection model. Created with BioRender.com. (B) Identification of infectivity genes. Plot shows rank-ordered mutants based on relative *in vivo* fitness scores. Dashed lines indicate relative fitness scores of −2.0 and 2.0 that were used to define phenotypes. Depleted or enriched mutants are highlighted. (C) Comparison of infectivity genes identified in the KN99α strain in C57BL/6J mice by KO-seq (this study) with those identified in the H99C/CM018 strain in A/J mice by signature-tagged mutagenesis^63^. (D) Repooling and retesting of infectivity mutants. Infectivity mutants from the primary screen were divided into 2 separate pools containing a constant set of 96 neutral mutants and were retested for *in vivo* fitness. Dashed lines indicate relative fitness scores of −2.0 and 2.0 that were used to define phenotypes. (E) Conservation levels and predicted domains of proteins encoded by 574 infectivity genes (left) and the 42 infectivity genes that lacked *in vitro* phenotypes (right). Outer circle represents the most specific conservation level for a protein based on OrthoMCL orthology groups. Inner circle indicates the presence or absence of bioinformatically predicted protein domains. (F) Overview of machine-learning based prediction of *in vitro* phenotypes predictive of *in vivo* fitness. (G) Normalized confusion matrix displaying average accuracy of model performance. (H) Feature importance (mean decrease in impurity; MDI) for the top 25 *in vitro* conditions predictive of *in vivo* fitness. Error bars represent standard deviation. Red bar represents median MDI across all 157 *in vitro* growth experiments.

We next sought to define host-relevant *in vitro* conditions. To this end, we trained a Random Forest Classifier to predict genes required for infectivity based on our *in vitro* phenotype data (Fig. 4F). We then performed a feature importance analysis to determine which *in vitro* conditions drove the predictive power of this classifier. We found that, as a group, genes required for infectivity could be predicted with 84% accuracy (Fig. 4G), indicating that the phenotype of most mutants with a fitness defect in the host could be explained by a requirement for growth under a particular environment. Feature importance analysis yielded a ranked list, of which the top 25 contributed significantly to sensitivity of the model (Fig. 4H, S6F, Table S6). Four of the top five most predictive environments correspond to tissue culture conditions (DMEM media, 37°C), supporting the relevance of this *in vitro* condition to infection. Other stress conditions and drug treatments were identified (Fig. 4H), indicating that host colonization specifically requires genes that enable tolerance to high salt and neutral/high pH, as well as cell wall integrity and ergosterol production.

Notably, cell wall synthesis and ergosterol production are targets for the two major classes of antifungal drugs, echinocandins and azoles^122^. The feature importance data leads us to propose that the efficacy of these drugs is related not only to their impact on viability, but also to the selective sensitivity of the host environment to inhibition of the pathways they target. For the azoles, this may also be related to hypoxia in the host, which limits oxygen-dependent ergosterol production regulated by the fungal SREBP pathway^49,50,52,123,124^.

### Tetraspanin-containing hyaluronic acid synthase complex promotes neurovirulence

Unlike those of model yeasts, the *C. neoformans* genome encodes a hyaluronic acid (HA) synthase, Cps1. Cps1 in the sister species *C. deneoformans* has been shown to promote invasion of the brain via the production of HA^125,126^. In *C. neoformans*, we observed that *cps1*Δ is not required for fitness in the mouse lung (Table S6) but does phenotypically cluster with deletions of two other genes, which, like Cps1, encode predicted transmembrane proteins: the fungal tetraspanin Tsp2 and a hypothetical protein encoded by *CNAG_05036,* which we refer to below as Cps2 (Fig. 5A, 5B). While tetraspanins have been described in fungi, they are thought to have a more primitive role in membrane deformation, and there is no evidence that they complex with other membrane proteins as they do in animal cells^127–130^. Nonetheless, we tagged Cps1 with an mNG-2XFLAG tag and performed AP-MS experiments on a detergent-solubilized membrane pellet. The top three most enriched proteins were Cps1, Tsp2, and Cps2 (Fig. 5C, Table S5). Consistent with the formation of a membrane complex, AlphaFold2-multimer predicts a three-protein complex with remarkably high confidence (Fig. 5D, Fig. S7A). Spinning-disk confocal microscopy revealed that Cps1-mNG-2XFLAG localizes to the cell surface (Fig 5E). In contrast, in *tsp2*Δ and *cps2*Δ strains, Cps1 displays a pattern characteristic of ER-localized proteins (Fig. 5E); this role for Tsp2 is reminiscent of the role of the CD81 tetraspanin, which is required for trafficking of CD19 B-cell coreceptor in mammals^131^. We tested whether Tsp2 and Cps2 were required for HA production. HA accumulation is nearly abolished in *cps1*Δ, *tsp2*Δ, and *cps2*Δ strains, a phenotype that could be complemented by expression of the corresponding wild-type alleles (Fig. 5F). These data demonstrate that a tetraspanin-containing complex is required for the assembly and trafficking of a hyaluronic acid synthase by *Cryptococcus neoformans*. This is the first demonstration of a tetraspanin-containing protein complex described outside animal cells. Whether and how it promotes the enzymatic function of Cps1 to utilize (presumably) cytosolic UDP-sugars and synthesize lumenal HA chains remains to be determined.

**Figure 5.**
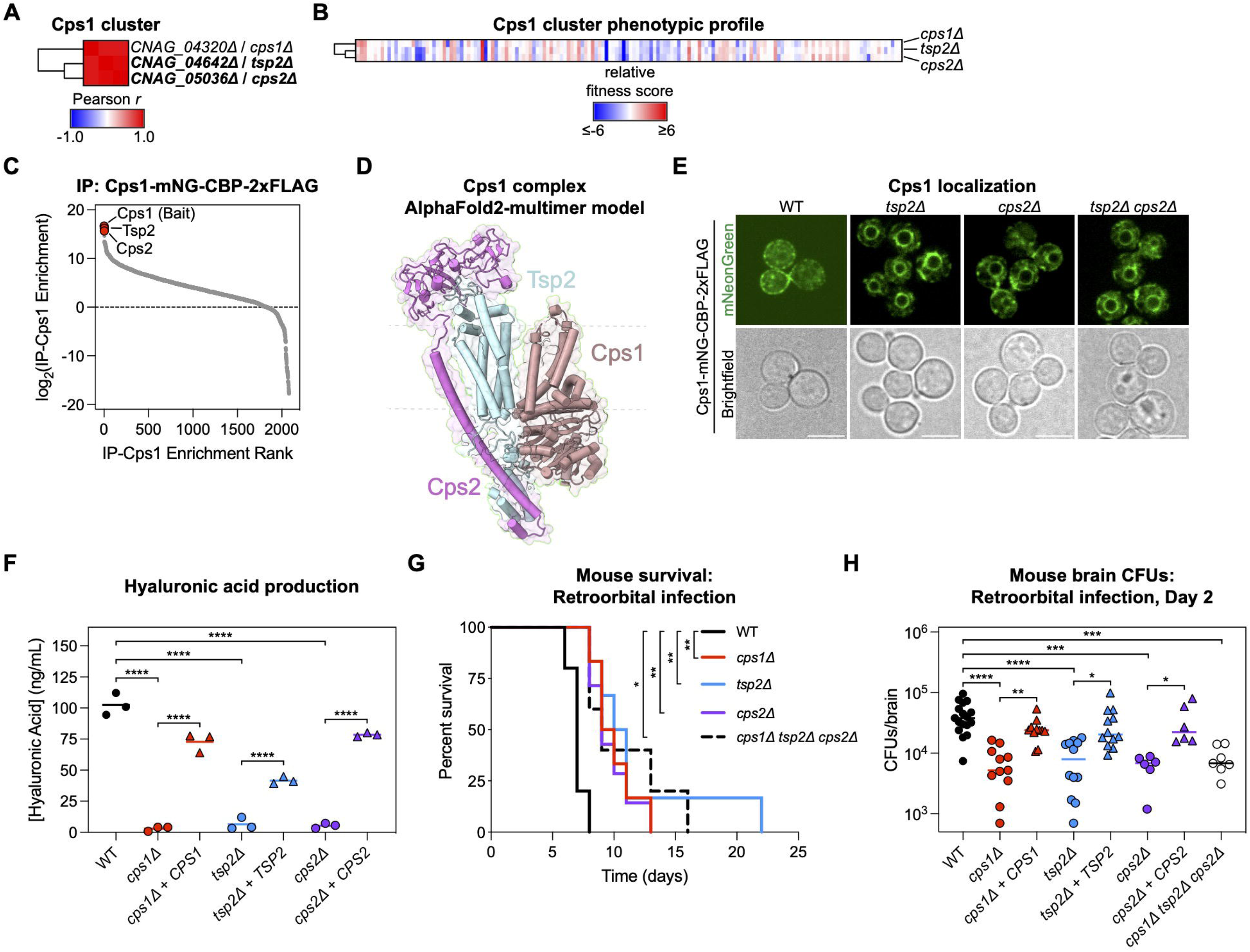
Tetraspanin-containing hyaluronic acid synthase complex promotes neurovirulence. (A) Cps1 cluster. (B) Phenotypic profile of Cps1 cluster members. (C) Cps1 AP-MS. Anti-FLAG AP-MS was performed on membrane extracts from a strain expressing endogenously tagged Cps1-mNG-CBP-2xFLAG. Data represent the protein-level ratio of normalized MS1 area from tagged versus untagged strains grown and processed in parallel. Averaged from 2 biological replicates. (D) AlphaFold2-multimer model of Cps1 cluster members. See also Figure S7A. (E) Live confocal imaging of endogenously tagged Cps1-mNG-CBP-2xFLAG. (F) ELISA assaying hyaluronic acid production from Cps1 cluster mutants. Bars represent mean values. (G) Survival of C57BL/6J mice infected intravenously (retroorbital inoculation) with 5 x 10^4^ CFUs of the indicated strains (*n* = 5-6 mice per group from 1 experiment). (H) Total brain CFUs 2 days post infection from mice infected intravenously (retroorbital inoculation) with 5 x 10^4^ CFUs of the indicated strains. Bars represent median values. WT, *n* = 17 mice pooled from 4 independent experiments with 3-6 mice per experiment; *cps1Δ* and *cps1Δ* + *CPS1*, *n* = 11 mice per group pooled from 3 independent experiments with 3-6 mice per experiment; *tsp2Δ* and *tsp2Δ* + *TSP2*, *n* = 12 mice per group pooled from 3 independent experiments with 3-6 mice per experiment; *cps2Δ*, *cps2Δ* + *CPS2*, and *cps1Δ tsp2Δ cps2Δ*, *n* = 6 mice per group pooled from 2 independent experiments with 3 mice per experiment. See also Figure S7B-D. Statistical analyses were performed with the Mantel-Cox test with Bonferroni correction for multiple hypotheses (G) and Kruskal-Wallis test with Dunn’s multiple comparisons test (H). \**p* < 0.05, \*\**p* < 0.01, \*\*\**p* < 0.001, \*\*\*\**p* < 0.0001.

Because Cps1 had been implicated in brain invasion in *C. deneoformans*^132,133^, we infected C57BL/6J mice with wild-type and mutant *C. neoformans* strains lacking *CPS1*, *TSP2*, or *CPS2* via retroorbital intravenous injection, which leads to rapid brain invasion. Mice infected with wild-type *C. neoformans* succumbed by day eight, but mice infected with knockout strains survived significantly longer (Fig. 5G). We measured brain and lung pathogen burden (colony forming units or CFUs) two and five days after infection (Fig. 5H, S7B-D). On day two after infection, mutant strains displayed significantly reduced brain CFUs relative to wild-type (Fig. 5H). This phenotype was rescued in the complemented strains (Fig 5H). We observed no difference in lung colonization between any strains tested at this time point (Fig. S7B), indicating a brain-specific phenotype. At five days post-infection, we observed no differences between the different strains in brain or lung CFUs (Fig. S7C, D), indicating an early kinetic role for HA production in brain colonization in *C. neoformans*, but not a complete block. This phenotype is not as dramatic as reported in *C. deneoformans*^125,126^, indicating that additional, partially redundant mechanisms promote neuroinvasion in *C. neoformans*.

### Inducible virulence factor required for adaptation to host-like conditions

Because fitness in tissue culture predicted genes required for infectivity (Fig. 4H), we sought genes with specific growth defects in this condition (DMEM, pH 7.4, 37°C, 5% CO_2_). This analysis identified a single gene, *CNAG_05159*, an unannotated cysteine-rich 16.5 kDa protein harboring a predicted *N*-terminal signal sequence (Fig. 6A, B) required for infectivity (Table S6). *CNAG_05159* was previously shown to be induced in tissue culture and in yeast isolated from the cerebrospinal fluid of an infected patient and is the direct target of several transcription factors that control virulence^134–137^. We confirmed this induction by quantitative reverse transcription PCR (Fig. 6C). We named the gene *RGH1* for *R*equired for *G*rowth Under *H*ost-like conditions. Using the knockout and complemented strains, we found that Rgh1 is required for virulence (Fig. 6D) and for accumulation of yeast in the lungs and brain of infected mice (Fig. 6E, F). An independent deletion strain constructed with CRISPR/Cas9 recapitulated these phenotypes (Fig. S8A-C). *RGH1* is required for growth under tissue culture conditions (Fig. 6A). It becomes dispensable, however, when the media, pH, temperature, or CO_2_ stresses that comprise “tissue culture conditions” are independently moderated. For example, at 30°C or pH 7.0, *RGH1* is dispensable for growth (Fig. 6G). This indicates that *RGH1* is required specifically under the combination of conditions that enable animal cells to grow in culture.

**Figure 6.**
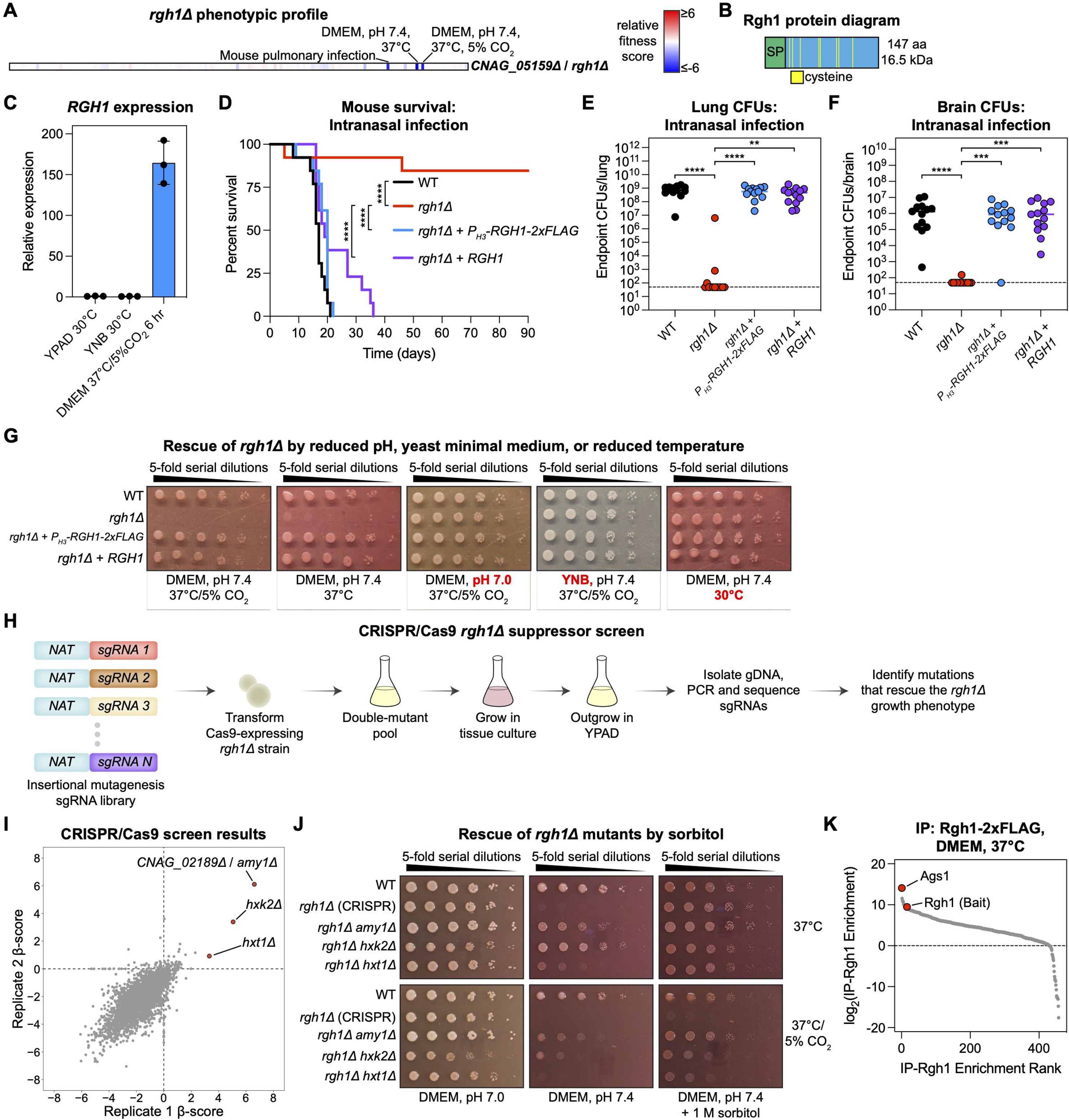
Inducible virulence factor required for adaptation to host-like conditions. (A) Phenotypic profile of the *rgh1Δ* mutant. (B) Rgh1 protein diagram. (C) qRT-PCR of *RGH1* transcripts in yeast cultured in standard laboratory conditions (YPAD or YNB at 30°C) or tissue culture (DMEM at 37°C/5% CO_2_). Expression is relative to the YPAD condition. Error bars represent standard deviation. (D) Survival of C57BL/6J mice infected intranasally with 5 x 10^4^ CFUs of the indicated strains (*n* = 13 mice per group pooled from 3 experiments with 3-4 mice per experiment). (E and F) Total endpoint CFUs isolated from the lungs (E) and brains (F) of mice in (D). Bars represent median values; dashed line represents limit of detection. Mice with undetectable CFUs were plotted at the limit of detection. (G) Spot dilution assays assessing growth of the indicated strains in tissue culture conditions (DMEM, pH 7.4, 37°C/5% CO_2_) and when CO_2_, pH, growth medium, or temperature stresses were removed. Images were taken after 2 days of growth. (H) Schematic for *rgh1Δ* suppressor screen using a genome-wide CRISPR/Cas9 insertional mutagenesis approach. (I) Suppressor screen results. Graph compares gene-level β-scores (fitness) from two independent sgRNA library transformations grown as shown in (H). (J) Spot dilution assays assessing growth of the indicated strains in tissue culture conditions in the absence or presence of 1 M sorbitol. Images were taken after 3 days of growth. (K) Rgh1 AP-MS. Anti-FLAG AP-MS was performed on clarified cell lysates from a strain expressing endogenously tagged Rgh1-2xFLAG grown in DMEM, pH 7.4 at 37°C. Data represent the protein-level ratio of normalized MS1 area from tagged versus untagged strains grown and processed in parallel. Averaged from 2 biological replicates. Statistical analyses were performed with the Mantel-Cox test with Bonferroni correction for multiple hypotheses (C) and Kruskal-Wallis test with Dunn’s multiple comparisons test (D and E). \**p* < 0.05, \*\**p* < 0.01, \*\*\**p* < 0.001, \*\*\*\**p* < 0.0001.

To gain insight into the function of Rgh1, we selected suppressors of the *rgh1*Δ growth defect under tissue culture conditions using a whole-genome CRISPR/Cas9-based insertional mutagenesis system we have developed (M. Huang et al., in preparation) (Fig. 6H). sgRNA sequencing identified a strong suppressor insertion in a predicted secreted α-amylase (*AMY1*) and weaker suppressors corresponding to insertions in *HXK2* (encoding hexokinase) and *HXT1* (encoding a hexose transporter) (Fig. 6I, Table S7). We confirmed these phenotypes by independently constructing double mutants (Fig. S8D).

Fungal amylases hydrolyze α-1,4-glucan linkages^138–141^, which are found in the outer layer of fungal cell walls as part of α-glucan, a critical outer cell wall component found in most fungi, including *C. neoformans*^142–144^. In *Aspergillus oryzae*, a related α-amylase acts on α-glucan to reduce its molecular weight, cleaving spacer α-1,4 linked glucan that interrupts α-1,3-glycan chains^145^. Thus, a plausible explanation for the suppression of the growth defect of *rgh1*Δ by *amy1*Δ is that *RGH1* promotes adaptive α-glucan production under stress conditions. To begin to test this model, we assessed whether Rgh1 impacts cell wall integrity under stress conditions. Indeed, we found that adding the osmotic stabilizer sorbitol to the media (known to suppress cell wall defects) can suppress the growth defect of *rgh1*Δ cells in DMEM at 37°C, although only weakly when 5% CO_2_ stress is also present (Fig. 6J). The enhanced ability of sorbitol to suppress the phenotype of *rgh1*Δ cells in *hxk*2Δ and *hxt1*Δ mutants suggests that additional mechanisms may be involved in the function of Rgh1 (Fig. 6I).

To gain insight into how Rgh1 may enhance cell wall integrity, we performed AP-MS experiments using extracts from cells grown in DMEM, pH 7.4 at 37°C. The co-purifying protein with the greatest signal (MS1 area) was the *C. neoformans* α-glucan synthase Ags1^143^ (Fig. 6K, Table S5). Given this biochemical observation and the genetic suppression of the growth defect of *rgh1*Δ by loss of a predicted cell wall α-amylase, we infer that Rgh1 promotes the production of α-glucan, potentially by directly activating Ags1. In summary, these data demonstrate the identification of a factor, Rgh1, induced under host-like conditions that enhances cell wall integrity, likely by regulating α-glucan production, a critical component of fungal cell walls. Together with prior work^143,144,146^, these findings point to α-glucan as a major vulnerability under infection conditions.

### Pathogen secreted factors promote infectivity

Secreted effectors underpin the virulence of most bacterial pathogens. A similar picture is beginning to emerge in fungi, but only a handful have been identified^147^. Because previous efforts at defining secreted proteins in *C. neoformans* did not consider lysed cells as a source for proteins identified in the culture supernatant^148–151^, we performed proteomic analysis of paired culture supernatants and cell pellets from *C. neoformans* grown in both laboratory media and under tissue culture conditions, optimizing growth conditions and lysis methods (Fig. 7A). We scored *bona fide* secreted proteins by ranking the quotient of protein signal in culture supernatants versus the cell pellet and then using a principled cutoff based on the maximum χ^2^ value produced by all possible cutoffs (Fig. 7B,C, Table S8). This analysis identified 66 high-confidence secreted proteins, 48 under each condition with an overlap of 30 proteins (Fig. 7D). These are highly enriched for proteins with predicted *N*-terminal signal sequences and/or internal hydrophobic segments that could serve as internal secretion signals (Fig. 7E).

**Figure 7.**
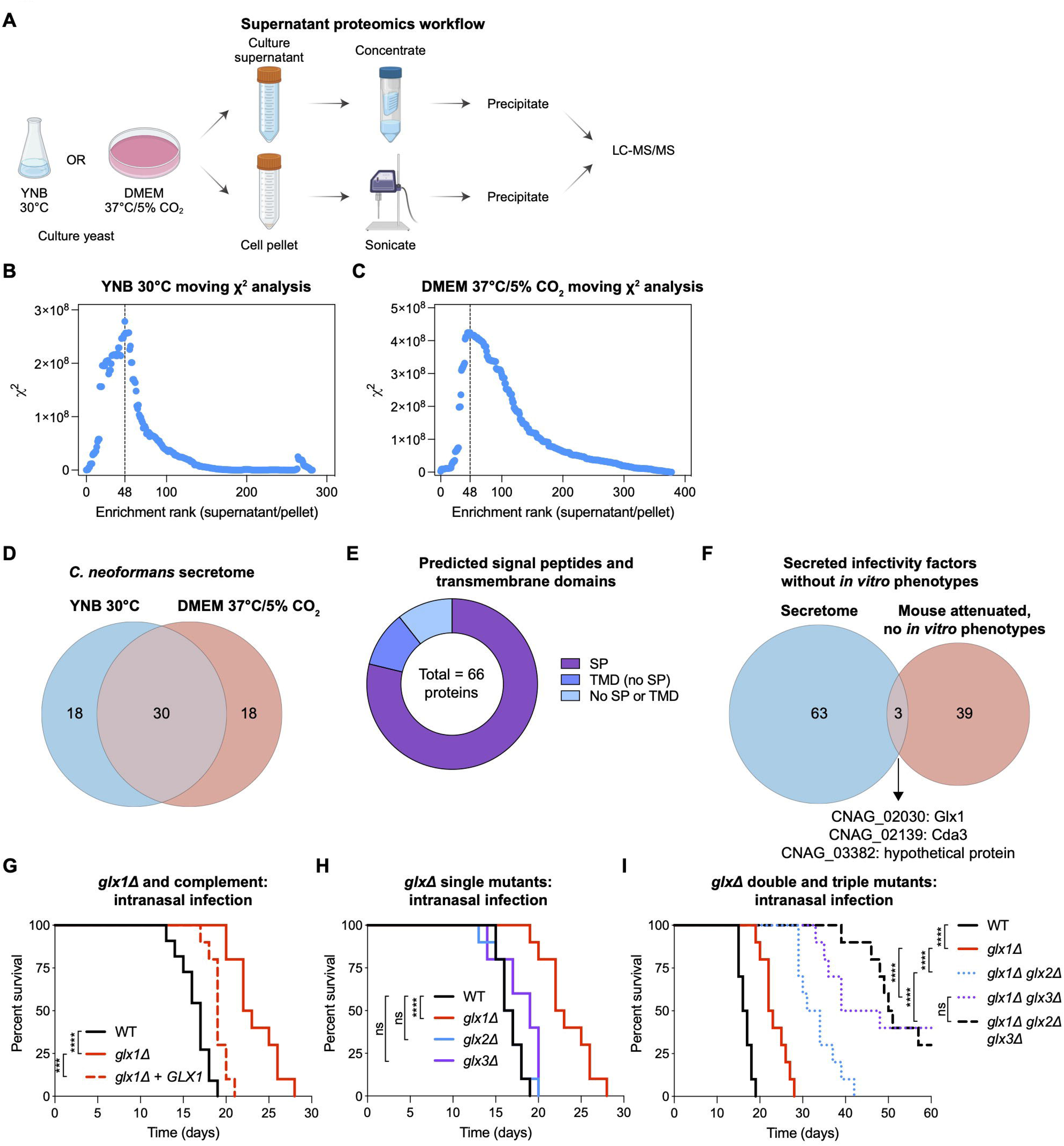
Pathogen secreted proteins required for infectivity. (A) Schematic of *C. neoformans* supernatant proteomics. Created with BioRender.com. (B, C) Moving χ^2^ analysis of proteins rank-ordered by enrichment in culture supernatants versus cell pellets in YNB at 30°C (B) or DMEM at 37°C/5% CO_2_ (C). Vertical dashed lines indicate maximum χ^2^ values used to set cutoffs for classifying proteins as secreted. (D) Overlap of secreted proteins identified from cultures grown in standard yeast growth conditions (YNB at 30°C) or in tissue culture (DMEM). (E) Predicted signal peptides and transmembrane domains in secreted proteins. (F) Overlap of secreted proteins identified with infectivity genes lacking *in vitro* phenotypes. (G-I) Survival of C57BL/6J mice infected intranasally with 5 x 10^4^ CFUs of the indicated strains. (G) *n* = 10-11 mice per group pooled from 2 experiments with 5-6 mice per experiment. (H) *n* = 10 mice per group pooled from 2 experiments with 5 mice per experiment. (I) *n* = 10 mice per group pooled from 2 experiments with 5 mice per experiment. Statistical analyses were performed with the Mantel-Cox test with Bonferroni correction for multiple hypotheses (G-I). ns, not significant, \*\*\**p* < 0.001, \*\*\*\**p* < 0.0001.

Among these, six displayed a defect in fitness in the mouse. Of these, three did not display an *in vitro* defect under the 141 conditions tested (Fig. 7F). Because they display a fitness defect in a pooled screen, the encoded proteins are likely to act locally. One corresponds to *CNAG_02030*, encoding a predicted copper-radical glyoxal oxidase we name Glx1. It is part of a family of three proteins in *C. neoformans* that also includes CNAG_00407 (Glx2) and CNAG_5731 (Glx3). In other fungi, all biochemically characterized members of this extracellular enzyme family (CAZy ID:AA5_1) use molecular oxygen to oxidize aldehydes, producing the corresponding carboxylic acid and hydrogen peroxide^152^. The *C. neoformans* Glx protein have the conserved active site residues (Fig. S8E). In rot fungi, this activity enables H_2_O_2_-requiring lignin peroxidases to break down lignocellulose, a major plant-derived nutrient source for fungi in the environment^153^.

Infection of mice revealed that a *glx1*Δ mutant is indeed attenuated, a phenotype that can be complemented by the wild-type allele (Fig. 7G). Single *glx2*Δ and *glx3*Δ mutants display no virulence defect (Fig. 7H), but their deletion in the *glx1*Δ background significantly enhanced the *glx1*Δ virulence defect, particularly for the *glx1*Δ *glx3*Δ double mutant, indicating partial redundancy (Fig. 7I). *In vitro*, the single- and double-mutants did not display defects in growth at 37°C in either laboratory or tissue culture media (Fig. S7F). Because all known members of the family to which Glx1-3 belongs are aldehyde-consuming, H_2_O_2_-producing proteins, we speculate that H_2_O_2_ production, aldehyde consumption, or production of an organic acid may promote virulence. These findings suggest that the Glx proteins, which likely have been selected for in the environment for saprotrophic purposes, are repurposed in the context of mammalian infection.

### Perspective

This work represents a functional analysis of the genome of a human meningitis fungal pathogen that lies at the top category of the W.H.O. priority fungal pathogens list. While constructing an ordered knockout collection represents a major investment of time and effort, it provides an impactful resource. Prepublication sharing of this strain library has provided other laboratories with a source of rigorously validated individual strains of interest and enabled several genome-wide screens^144,154–171^. Before this work, knockout collections that included the majority of nonessential genes were publicly available for three fungal species, all non-pathogenic ascomycetes (*S. cerevisiae*, *S. pombe*, and *Neurospora crassa*). These were constructed by either a consortium of academic laboratories or a for-profit entity^172–174^. To demonstrate the utility of the *C. neoformans* resource, we used pooled approaches and a high-precision sequencing-based method to connect genotype to phenotype at scale, obtaining over a million high-quality measurements.

We have shown how these data can be exploited as hypothesis-generation tools to drive experimental work producing insights into *C. neoformans* biology. Although *S. cerevisiae* has long dominated fundamental studies of cell biology, many important pathways continue to be deciphered in other unicellular systems, including *S. pombe* and *N. crassa*, and recent systematic studies of *C. reinhardii* have shown promise for understanding photosynthesis^27,28^. Our discovery of a UNC79/UNC80 pair in *C. neoformans* and an NHEJ-promoting activity of its 53BP1 ortholog revises current views on the evolution of these pathways. It also points to the value of supporting fundamental research beyond a small set of model organisms.

We have also described how the knockout collection and phenotypic atlas can be used to advance our understanding of infection biology. Our work shows how integrating unbiased measurements of *in vitro* and *in vivo* phenotypes with biochemical and cell biological analysis can lead to insights that would be difficult or impossible to obtain through piecemeal investigations. Analysis of genes required for fitness in specific environments and their relationship to those required in the host illuminate environmental adaptations that enable host colonization. Likewise, analysis of pathogen genes required for fitness in the host, but without a detectable *in vitro* defect, points to unanticipated mechanisms for virulence.

### Limitations of this study

The knockout collection we constructed covers approximately 80% of nonessential genes. Further efforts will be required to construct mutations in the remaining nonessential gene set. Essential genes are not analyzed here, yet hypomorphic mutations are valuable for understanding cellular pathways^175^. We have recently established the auxin degron system in *C. neoformans* (M. Huang et al., in preparation), which will enable the generation of such mutants. We note that profiling the library under additional environmental conditions might yield phenotypes beyond the 1909 genes for which we obtained phenotypes. In addition, different strain backgrounds may yield distinct genotype-phenotype relationships. Our pooled infection strategy may miss factors that act non-locally and whose loss can, therefore, be complemented by other mutants in a pool. Because the gene deletion strains have not been whole-genome-sequenced, unlinked mutations (which are rare^176^) could be responsible for a given phenotype, necessitating the regeneration and/or complementation of mutants to confirm genotype-phenotype relationships. Likewise, many of the gene clusters identified have yet to be validated and investigated mechanistically. Finally, purification of active recombinant enzymes will be required to demonstrate that the *C. neoformans* Glx proteins have H_2_O_2_-producing activity.

## MATERIALS AND METHODS

### Microbial strains and growth conditions

*C. neoformans* strains used in this study are listed in Table S9. Yeast were cultured in YPAD, YNB, low-iron YNB (LI-YNB), synthetic complete (SC), or DMEM as indicated. YPAD contained 1% yeast extract (Gibco 212720), 2% peptone (Gibco 211820), 2% glucose, 0.15 g/L L-tryptophan, and 0.04 g/L adenine. YNB contained 1.5 g/L yeast nitrogen base powder without ammonium sulfate (MP Biomedicals 4027032), 5 g/L ammonium sulfate, and 2% glucose. LI-YNB contained 6.9 g/L YNB with ammonium sulfate and without iron (ForMedium CYN1101) and 2% glucose. SC contained YNB with 22 mg/L uracil, 21 mg/L adenine, 8.6 mg/L para-aminobenzoic acid, 174 mg/L L-leucine, 80 mg/L L-tryptophan, and 85 mg/L of each L-alanine, L-arginine-HCl, L-asparagine, L-aspartic acid, L-cysteine-HCl, glutamine, L-glutamic acid, glycine, myo-inositol, L-isoleucine, L-methionine, L-phenylalanine, L-proline, L-serine, L-threonine, L-tyrosine, L-valine, L-histidine-HCl, and L-lysine-HCl. For all experiments except *RGH1* qRT-PCR and supernatant proteomics, DMEM contained 13.48 g/L DMEM powder without sodium bicarbonate (Gibco 12800082), 50 U/mL penicillin, 50 μg/mL streptomycin (PenStrep; Gibco 15070063), 2 mM additional L-glutamine (Corning 25-005-CI), and 100 mM HEPES, adjusted to desired pH with NaOH. For *RGH1* qRT-PCR, DMEM with sodium bicarbonate (Corning 10-013-CV) was supplemented with 10 mM HEPES (Corning 25-060-CI) plus PenStrep and L-glutamine as above. For supernatant proteomics, phenol red-free DMEM with sodium bicarbonate (Corning 17-205-CV) was supplemented with 10 mM HEPES, PenStrep, and L-glutamine as above. Media buffered with McIlvaine’s buffer contained 77.1 mM Na_2_HPO_4_ and 61.5 mM citric acid (pH 4.0); 126.3 mM Na_2_HPO_4_ and 36.9 mM citric acid (pH 6.0); 164.7 mM Na_2_HPO_4_ and 17.7 mM citric acid (pH 7.0); or 194.5 mM Na_2_HPO_4_ and 2.8 mM citric acid (pH 8.0). For solid media, 2% agar was added. Routine culture was performed at 30°C unless otherwise specified. Liquid culture was performed on an orbital shaker (200 rpm) or roller drum. For genetic manipulations, strains were selected on agar plates containing 125 μg/mL nourseothricin sulfate (RPI N51200 or Jena Biosciences AB0-102XL), 300 μg/mL Hygromycin B (Calbiochem 400050), or 200 μg/mL G418 sulfate (Corning 61-234-RG).

### Molecular cloning

Molecular cloning was performed using HiFi DNA Assembly Master Mix (NEB E2521L). Plasmids and their corresponding backbones, inserts, and primers used for construction are listed in Table S9.

### Knockout library construction

Knockout library construction was performed with biolistic transformation as described previously^177^. Deletion constructs were purchased from Radiant Genomics, Inc. and consisted of a NAT resistance cassette flanked by ∼1 kb homology arms corresponding to the regions immediately upstream and downstream of the target gene’s predicted coding sequence (CDS). Genomic boundaries of genes targeted for deletion are listed in Table S1. Constructs were transformed by biolistic transformation and were selected on YPAD + NAT plates at room temperature for 3 days. From each transformation, at least 12 colonies were assessed for the expected knockout junctions and loss of CDS by colony PCR. We attempted deletions of each gene at least twice. This yielded 4692 strains. A second round of PCR quality control was performed with a set of independent “ORF-check” primers with a smaller amplicon size (∼0.5 kb vs. ∼1 kb) to assess a second time the presence or absence of the target CDS in knockout strains, identifying 291 false positives. Finally, during these studies, we identified strains that had diplodized, based on an unexpected shared phenotypic signature of functionally unrelated gene deletions followed by confirmation using DNA content measurements.

### Strain construction with CRISPR/Cas9

*C. neoformans* strains were constructed using CRISPR/Cas9 as previously described^76^. Primers and plasmids are listed in Table S9. Briefly, for each transformation we used 700 ng sgRNA, 1 μg *CnoCAS9* expression cassette, and 2 μg donor DNA with homology arms. All components were generated by PCR, except for donor DNA for Save Haven 1 (SH1) transformations, which was produced by linearization of plasmid DNA with AscI or BaeI as indicated in Table S9. To construct sgRNAs with unique 20 bp spacers, we first amplified separate “U6 promoter-unique spacer” fragments (MJB537 paired with guide-specific sgRNA_R primer) and “unique spacer-scaffold-6T terminator” fragments (guide-specific sgRNA_F primer paired with MJB538) from pBHM2329. We then mixed equal volumes of these products and performed fusion PCR with MJB535 and MJB536. The *CnoCAS9* expression cassette was amplified from pBHM2403 with MJB537 and MJB538. For gene deletion, *NAT*, *NEO*, or *HYG* expression cassettes with homology arms (typically 50 bp) were amplified from pSDMA25, pSDMA57, or pSDMA58, respectively. For endogenous *C*-terminal gene tagging, the desired tag-marker combination was PCR amplified from pBHM2404, pBHM2406, or pBHM2502.

For transformation, overnight cultures grown in YPAD were diluted to an OD_600_ of 0.2 in 100 mL YPAD and cultured 4-5 hours to an OD_600_ of 0.8-1.0. Yeast were pelleted at 3000 x*g* at 4°C and washed twice with 25 mL of cold water. Pellets were resuspended in 10 mL electroporation buffer (10 mM Tris-HCl, pH 7.4, 1 mM MgCl_2_, 270 mM sucrose) with 1 mM DTT and incubated on ice for 1 hr. Samples were then pelleted at 3000 x*g* at 4°C and resuspended in 250 μL electroporation buffer without DTT. For each transformation, 45 μL of resuspended cells were mixed with DNA, transferred to a prechilled 0.2 cm electroporation cuvette, and electroporated at 500 V, 400 Ω, and 250 μF on a BTX Gemini X2 electroporator. Yeast were resuspended in 1 mL YPAD and incubated rotating at 30°C for 2 hr before plating on selective media. Successful transformants were validated by diagnostic colony PCR^76^.

### Knockout library pooling

The *C. neoformans* deletion library contains 4692 strains arrayed into 49 96-well plates. Strains were grown as patches in 96-well arrayed format on YPAD + NAT agar in Nunc OmniTrays. Mutants within each plate were pooled by gently scraping yeast from the arrayed agar plates, washing the plate with YPAD to collect remaining cells, and resuspending in 15 mL final volume YPAD. We measured the OD_600_ of each of these 49 subpools and mixed equal OD_600_’s of each to generate either a master pool containing mutants from all 49 plates (for *in vitro* library profiling experiments) or 10 nonoverlapping subpools containing mutants from up to 5 plates (for *in vivo* experiments). Glycerol was added to each pool to 25% final concentration, and aliquots of 10 OD_600_’s each were stored at −70°C.

### *In vitro* fitness profiling

To profile mutant fitness *in vitro*, a frozen aliquot of the pooled deletion library was thawed, inoculated into 100 mL YPAD, and recovered overnight shaking at 200 rpm at 30°C. Cultures were washed once in either PBS or the base medium for the desired growth condition (typically YNB) and inoculated at an OD_600_ of 0.01 in 100 mL in the condition of interest and a corresponding control condition. All cultures were grown in duplicate. For most experiments, cultures were grown for 24 hr in YNB at 30°C in the presence of a chemical stressor and were compared to an untreated control. Compound concentrations were chosen empirically with the goal of stressing cultures sufficiently to identify mutants with differential fitness while minimizing population bottlenecks. We typically accomplished this with concentrations that inhibited the terminal OD_600_ of the culture by 20-60% compared to the untreated control. When we could not achieve this inhibition due to low efficacy, compound solubility limits, or excessively large quantities of required compound, we treated at the highest practical concentration. At the end of the growth period, cultures were harvested, washed once with water, and stored at −70°C until subsequent gDNA isolation and KO-seq library preparation.

In some experiments, we varied growth medium (e.g., YNB, LI-YNB, YPAD, or DMEM), pH (unbuffered, 4.0, 6.5, 7.0, 7.4, or 8.0) incubator conditions (30°C, 37°C, or 37°C/5% CO_2_), or carbon sources. Some of these conditions required adjustments to the standard parameters described above. For example, in a subset of growth conditions, particularly those involving DMEM, high temperature, and/or high pH, we found that our standard extraction protocol failed to yield sufficient gDNA for KO-seq, presumably due to difficulty in lysing yeast grown in highly stressful conditions. To circumvent this challenge, at the end of the growth period we pelleted cultures, washed once in YPAD, and inoculated 100 mL YPAD cultures at an initial OD_600_ of 0.1. We then grew these cultures overnight 30°C and harvested them for gDNA isolation and KO-seq as above.

For alternative carbon source experiments, we omitted glucose from YNB and added 2-3% of the desired carbon source. For a subset of these conditions, as well as for cultures grown in at pH 8.0, yeast did not achieve sufficient density for gDNA isolation within the standard 24 hr growth timeframe. We therefore measured the OD_600_’s of these cultures every 24 hrs post inoculation and harvested when they reached an OD_600_ ≥ 3.0 or when it appeared that growth had saturated.

In another experiment, we depleted iron from cultures and assessed mutant fitness in low-iron medium supplemented with different iron sources. To deplete cultures of iron, we grew the deletion library for 3 days in LI-YNB supplemented with 100 mM MOPS, pH 7.0 and 100 μM of the iron chelator bathophenanthrolinedisulfonic acid (BPDS). We then inoculated these iron-depleted cultures into that same growth medium supplemented with different iron sources and cultured for 2 days. These cultures were then outgrown in YPAD overnight as described above.

Table S2 summarizes the conditions of each growth experiment, including growth medium, pH, incubator conditions, chemical stressors, culture time, OD_600_ at the end of the growth period, relevant control condition, percent growth relative to control, and whether a YPAD outgrowth was used.

### Mouse infections

All animal experiments were approved by the UCSF Institutional Animal Care and Use Committee (IACUC) under protocol numbers AN144517, AN180537, and AN195144. Wild-type C57BL/6J mice were purchased from Jackson Laboratories (strain number 0000664) and were either used directly for experiments or were bred in-house. Excess dectin-1 heterozygous (*Clec7a^+/-^*) mice were used for pooled retest experiments and were derived from *Clec7a^-/-^*mice (Jackson strain number 012337) that were bred in-house with wild-type C57BL/6J mice. *Clec7a^+/-^*mice behave like wild type with regards to *C. neoformans* infection.

To profile the knockout library in a mouse pulmonary infection model, frozen aliquots of library subpools (containing up to 480 mutants) were thawed, inoculated into 100 mL YPAD, and recovered overnight shaking at 200 rpm at 30°C. Cultures were washed twice in sterile saline, counted on a hemocytometer, and adjusted to 2 x 10^6^ yeast/mL. For each subpool, 5 6-8-week old female mice were anesthetized with 75 mg/kg ketamine and 0.5 mg/kg dexmedetomidine and suspended them by their front incisors from a silk thread. Mice were infected by pipetting 50 μL yeast suspension (1 x 10^5^ yeast total) into the nares and were kept suspended for 10 additional min, after which anesthesia was reversed with 1 mg/kg atipamezole. Infection was allowed to progress until mice reached a predefined experimental endpoint (15% weight loss relative to preinfection or other signs of severe disease), at which they were euthanized by CO_2_ inhalation followed by cervical dislocation. To isolate yeast for gDNA isolation, lung tissue was harvested, resuspended in 5 mL final volume PBS, homogenized, and cultured in 100 mL YPAD shaking at 200 rpm at 30°C overnight. Control cultures were generated by similarly culturing the inocula used to infect mice in YPAD overnight. Cell pellets were collected by centrifugation, washed in water, and stored at −70°C until later processing. In the primary screen, cultures from one mouse in Pool 8 and one mouse in Pool 9 were lost due to bacterial contamination. Fitness retest experiments performed in *Clec7a^+/-^* mice were performed similarly except that 1) each group of 5 mice contained 3 females and 2 males; 2) 14 days post infection was used for the experimental endpoint; and 3) lung homogenates were cultured in the presence of 50 μg/mL chloramphenicol to prevent bacterial contamination.

Monotypic intranasal infections were performed similarly to pooled screens except that we used both male and female mice aged 6-12 weeks (age- and sex-matched between experimental groups); mice were infected with 5 x 10^4^ yeast; and a weight loss threshold of 20% was used to define experimental endpoint. For retroorbital infections, yeast preparation and mouse analgesia were performed similarly, except that yeast inocula were prepared at 5 x 10^5^ CFU/mL and 100 μL were delivered by retroorbital injection to achieve a final inoculum of 5 x 10^4^ yeast. To enumerate CFUs from harvested lung or brain tissue, serial dilutions of homogenate were plated onto YPAD plates and grown for 2 days at 30°C. Statistical differences in survival and CFU counts were analyzed in GraphPad Prism (version 10.2.3).

### Genomic DNA (gDNA) isolation

gDNA for KO-seq was isolated as previously described^177^ with modifications. Frozen cell pellets from 100 mL of culture were lyophilized overnight followed by resuspension in 10-15 mL CTAB lysis buffer (100 mM Tris-HCl, pH 7.4, 700 mM NaCl, 10 mM EDTA, 1% CTAB, 1% β-mercaptoethanol) with 3-5 mL of 0.5 mm zirconia/silica beads (BioSpec 11079105z). Samples were vortexed and incubated in a 65°C water bath for 2 hours with intermittent vortexing. An equal volume of chloroform was added, and samples were gently mixed followed by centrifugation at 2000 x*g* for 20 min. The aqueous layer was collected, and DNA was precipitated with an equal volume of isopropanol, pelleted at 2000 x*g* for 10 min, washed with 5 mL 70% ethanol, and air-dried. DNA was resuspended in 500 μL 10 mM Tris, pH 8.0 overnight and was treated with 20 μg/mL RNase A (Thermo EN0531) at 37°C for 30 min followed by 200 μg/mL proteinase K at 55°C for 1 hour. An equal volume of phenol:chloroform:isoamyl alcohol (25:24:1) was added and the sample was vortexed and centrifuged at 21,000 x*g* for 10 min. The aqueous layer was removed and extracted with 500 μL chloroform, vortexed, and centrifuged at 21,000 x*g* for 10 min. The aqueous layer was removed, precipitated with 0.3 M sodium acetate and 3 volumes of ethanol, and centrifuged at 21,000 x*g* for 10 min. The pellet was washed with 70% ethanol, air-dried, and resuspended overnight in 100 μL 10 mM Tris-HCl, pH 8.0. DNA was further cleaned with a genomic DNA clean & concentrator-25 kit (Zymo D4065) and was quantified with the Qubit dsDNA broad range kit (ThermoFisher Q32853).

gDNA isolation for the CRISPR/Cas9 suppressor screen (see below) was performed similarly with modifications. Briefly, frozen cultures from initial mutant pools and each assay condition were treated with DNAse to remove any extracellular PCR product. 100 OD_600_’s of cells were treated with 60U DNase I (NEB M0303) and were incubated at 37°C for 1.5 hours followed by heat inactivation at 75°C for 30 minutes. Cells were washed with water, resuspended in 6 mL CTAB lysis buffer, and transferred to Omni 15 ml sample tubes (Omni International 19-6615) with approximately 1.5 mL of 0.5 mm zirconia/silica beads. The cells were bead-beaten for two rounds of 1.5 minutes at 5 m/s at 4°C in an Omni Bead Ruptor Elite (Omni International 19-042E), then incubated for at least 2 hours at 65°C. After incubation, 2 ml of 25:24:1 phenol:chloroform:isoamyl alcohol was added, and cells were bead-beaten for another round using the same settings. The aqueous phase was then collected by centrifugation, and DNA was precipitated by isopropanol precipitation as described above. DNA was rehydrated with 100 μL nuclease-free water and digested using 10 μg RNase A with incubation at 37°C for 30 minutes, followed by treatment with 100 μg Proteinase K with incubation at 55°C for 1 hour. Samples were purified using 500 μL of 25:24:1 phenol:chloroform:isoamyl alcohol extraction, then ethanol precipitated and washed in ethanol twice. The resulting pellet was then resuspended in 100 μL nuclease-free water and cleaned with a Zymo gDNA Clean and Concentrator column as described above.

### KO-seq library preparation and sequencing

Two separate library preparations, one for upstream KO junctions and one for downstream junctions, were performed for each gDNA sample using the NEBNext Ultra II FS kit (NEB E7805L) with modifications, including a custom Illumina adaptor and primers. The custom adaptor contains 1) an 8N unique molecular identifier (UMI) in place of the i7 index; 2) a truncated P5 branch ending with dideoxycytosine; and 3) an extension from the typical P7 primer binding site that enables nested PCR during library preparation. To create this adaptor, primers MJB002 and MJB049 were resuspended to 200 μM in 10 mM NaCl. Equal volumes were mixed, and primers were annealed by heating to 95°C for 5 min followed by cooling at a rate of 0.1°C/s to 12°C. Adaptors were diluted to 15 μM with water, aliquoted, at stored at −20°C.

To prepare sequencing libraries, 1 μg input gDNA was adjusted to 26 μL final volume with 10 mM Tris-HCl, pH 8.0. To fragment, end-repair, and A-tail gDNA, 7 μL Ultra II FS Reaction Buffer and 2 μL Ultra II FS Enzyme Mix were added, and samples were incubated at 37°C for 20 min followed by 65°C for 30 min. For adaptor ligation, 30 μL Ultra II Ligation Master Mix, 1 μL Ligation Enhancer, and 2.5 μL 15 μM preannealed adaptor were added, and samples were incubated at 20°C for 15 min. Samples were adjusted to 100 μL with 10 mM Tris-HCl, pH 8.0 and were dual size-selected with AMPure XP beads (Beckman A63881). To remove large DNA fragments, 0.4 volumes (40 μL) of beads were added. Samples were vortexed, incubated at room temperature for 10 min, and placed on a magnet stand for 5 min, and the supernatant was collected and transferred to a fresh tube. To remove small fragments, 0.2 volumes (20 μL) of beads were added to the supernatant. Samples were vortexed, incubated at room temperature for 10 min, and placed on a magnet stand for 5 min. The supernatant was removed and discarded, and beads were washed twice with 200 μL 80% ethanol and air-dried. DNA was eluted by addition of 16 μL distilled water, and 15 μL was transferred to a fresh tube.

For the first round of PCR, we used a forward primer that anneals to the NAT resistance cassette in each deletion strain (MJB037 for downstream junctions or MJB038 for upstream junctions) and a reverse primer that anneals to the extended P7 region of our custom adaptor (MJB050). 25 μL of Q5 Master Mix and 5 μL of each 10 μM primer was added to each 15 μL DNA sample, and PCR was performed with an initial denaturation of 98°C for 30 s, 12 cycles of 98°C for 10 s and 65°C for 75 s, and a final extension of 65°C for 5 min. To clean samples, 0.9 volumes (45 μL) of AMPure XP beads were added, and samples were vortexed, incubated at room temperature for 10 min, and placed on a magnet stand for 5 min. The supernatant was removed and discarded, and beads were washed twice with 200 μL 80% ethanol and air-dried. DNA was eluted by addition of 16 μL distilled water, and 15 μL was transferred to a fresh tube.

For the second round of PCR, we used forward primers that anneal internal to those used in the first PCR and a reverse primer (MJB039) that anneals to the canonical P7 region of the Illumina adaptor. Forward primers contained the complete Illumina P5 region and 8 bp sample barcodes in the i5 index site and are listed in Table S9. Additionally, forward primers contained a stagger that inserts 0-3 random nucleotides between the Illumina Read 1 primer binding site and the NAT cassette-annealing region. This stagger randomizes the first bases of Illumina Read 1, which would otherwise be identical for each molecule due to the constant NAT cassette-annealing region of this primer comprising the first bases of the read. To create this stagger, we mixed equimolar amounts of 4 otherwise identical primers containing 0-3 N’s in the desired region. This PCR was performed using the same conditions as the first PCR above and was similarly cleaned with AMPure XP beads, except that a 30 μL elution volume was used.

Library concentrations were measured on an Agilent 2100 Bioanalyzer using the high sensitivity DNA kit (Agilent 5067-4626) or TapeStation 4200 using the high sensitivity D1000 system (Agilent 5067-5584). Samples were pooled at the desired molar ratios and were sequenced with paired-end 100 bp reads on a HiSeq 4000 or NovaSeq 6000 at the UCSF Center for Advanced Technology (CAT) or with paired-end 75 bp reads on a NextSeq 500 at the Gladstone Genomics Core.

### KO-seq data analysis

Raw sequencing files were demultiplexed at the UCSF CAT using the i5 index (Index 2) added in the second round of KO-seq PCR. The i7 index (Index 1) contains an 8 bp UMI to be used later for read deduplication. UMI sequences corresponding to each read were appended to the sequence identifier lines of Read 1 and Read 2 FASTQ files with a custom Python (version 3.7.1) script.

Reads were trimmed and filtered with Cutadapt (version 2.4)^178^ In addition to standard adaptor removal, we used this step to remove reads resulting from mispriming during library preparation. Briefly, the first 22 bp of KO-seq Read 1 sequences are constant, consisting of the 16 bp annealing region of the PCR 2 forward primer followed by the final 6 bp of the NAT cassette. These sequences are CGGCCGCATCCCTGCATCCAAC for downstream junctions and CGCGCCTAGCAGCGGATCCAAC for upstream junctions. Reads that lack this full 22 bp sequence likely resulted from mispriming during PCR and were discarded. We used a quality cutoff of 20, a maximum error rate of 20%, and a minimum read length of 20 bp for this step.

The command-line calls for this step were

cutadapt -g “CGGCCGCATCCCTGCATCCAAC;required;o=22;e=0.2…AGATCGGAAGAGCACACGTCTGA ACTCCAGTCAC;optional” -A GTTGGATGCAGGGATGCGGCCG -q 20 -m 20 --pair-filter=first -- discard-untrimmed -o read1_out1 -p read2_out1 read1_fq read2_fq” for downstream junctions and

cutadapt -g “CGCGCCTAGCAGCGGATCCAAC;required;o=22;e=0.2…AGATCGGAAGAGCACACGTCTGAACTCCAGTCAC;optional” -A GTTGGATCCGCTGCTAGGCGCG -q 20 -m 20 --pair-filter=first -- discard-untrimmed -o read1_out1 -p read2_out1 read1_fq read2_fq for upstream junctions. We subsequently discarded any trimmed reads that were less than 20 bp in length using

cutadapt -m 20:20 -o read1_out2 -p read2_out2 read1_out1 read2_out1

Deletion mutants were constructed such that each gene’s CDS was replaced with the NAT cassette while leaving the endogenous UTRs in place. We therefore used Bowtie2 (version 2.3.5.1)^179^. To map trimmed reads to a custom genome index relating each annotated *C. neoformans* gene to the 300 bp immediately preceding or following the region deleted. We did this with the command line call

bowtie2 --end-to-end --fr --norc --no-mixed --no-discordant -X500 --score-min C,-22,0 -p threads -x index -1 read1_out2 -2 read2_out2 -S sam_name

The resulting SAM file was then converted to BAM format, sorted, and indexed using SAMtools^180^ implemented in pysam (version 0.15.3). Reads resulting from PCR duplicates were deduplicated based on the combination of their 8 bp UMI and the random mapping coordinate of KO-seq Read 2 using UMI-tools (version 1.0.1)^181^ with the command line call umi_tools dedup -- paired --output-stats=name --log=name.log --stdin=sorted_bam --stdout=deduplicated_bam The number of deduplicated reads corresponding to each gene flank were counted using a custom Python script, excluding reads that mapped >10 bp from the start of each gene flank. Counts corresponding to mutants that failed quality control or had diploidized (Table S1) were also discarded.

### *In vitro* mutant fitness calculation

Mutant fitness calculations were performed using custom scripts in R (version 4.3.2). Upstream and downstream junction counts were summed for each mutant in each sample and were used as input for DESeq2 (version 1.40.2)^182^ to calculate mutant log_2_ fold changes in conditions of interest relative to same-day controls. DESeq data sets were constructed using the DESeqDataSetfromMatrix function, and the DESeq function was called with the “fitType = ‘local’” parameter. Samples cultured on the same day (i.e., within the same experiment in Table S2) were analyzed as individual batches. Log_2_ fold changes were shrunk using the lfcShrink function implementing the ashr (version 2.2_63)^183^ shrinkage estimator. To compare log_2_ fold changes across conditions and experiments, we scaled ashr-shrunk log_2_ fold changes by dividing them by the standard deviation of all log_2_ fold changes within a sample and defined this value as a mutant’s “relative fitness score.”

### *In vivo* mutant fitness calculation

Upstream and downstream junction counts were summed for each mutant in each mouse and normalized by converting to reads per million with a pseudocount of 1. Only mutants for which we obtained at least 10 raw counts and 10 counts per million in the control YPAD culture were analyzed. Raw log_2_ fold changes for each mutant in each mouse were calculated relative to their abundances in the control YPAD culture of the infectious inoculum. Because mutants are more likely to display decreased rather than increased fitness *in vivo*, raw log_2_ fold changes exhibited asymmetrical lognormal distributions that were centered near zero, as expected, but displayed a long tail below zero. A small subset of samples were centered below zero, presumably due to noise. These features precluded use of the simple scaling approach described for *in vitro* data to correct for mouse-to-mouse variability. Instead, we transformed raw log_2_ fold changes to modified (median-based) *Z*-scores, *Z_m_*, and defined population outliers as mutants with |*Z_m_*| > 3.5. We calculated the mean and standard deviation of raw log_2_ fold changes for non-outlier mutants, then used these values to calculate standard *Z*-scores for all mutants, including outliers. We then averaged these values for each mutant across replicate mice to produce an *in vivo* relative fitness score for each mutant.

For retest experiments, we created 2 subpools containing 368 or 367 non-overlapping putatively attenuated mutants based on a preliminary analysis of the initial mouse screen. These subpools respectively contained 18 and 6 mutants later discovered to be diploid (Table S1), which were omitted from analyses. As controls, to each pool we added the same set of 96 neutral mutants, which had fitness scores near zero in the preliminary analysis. Our *in vivo* fitness calculations for the initial screen described above relied on the fact that most mutants in the initial pools had fitness similar to wild type, thereby allowing us to identify attenuated mutants as those that deviated from the population average. We could not apply this method to our retest experiment because retest pools primarily consisted of attenuated strains. Instead, for the retest experiment we first calculated raw log_2_ fold changes for each mutant (relative to the control culture of the infectious inoculum) as described for the initial screen. We then calculated the mean and standard deviation of these log_2_ fold changes for the 96 control mutants in each pool. Finally, we normalized the raw log_2_ fold change of each attenuated mutant using the mean and standard deviation of the control strains. We averaged these values from the 5 replicate mice to obtain a relative fitness score for each mutant. Because the 2 attenuated mutant pools were designed based on an early analysis of the primary screen data, they contained 53 and 87 mutants, respectively, that were no longer classified as “attenuated” after performing the more rigorous analysis described above. These mutants are excluded from the graph in Fig. 4D for clarity but are reported in Table S6.

### Hierarchical clustering

*In vitro* and *in vivo* mutant fitness scores across 158 growth experiments (157 *in vitro* and 1 *in vivo*) were compiled into a single data frame. We defined mutants as having a phenotype in a particular growth condition if the absolute value of their fitness score was greater than or equal to 2.0. We filtered this data frame to include only mutants with a phenotype in at least 1 experiment, creating a matrix of 1909 mutants in 158 experiments. This matrix was hierarchically clustered using average linkage clustering with the centered Pearson correlation as the distance metric in Cluster 3.0 (C Clustering Library version 1.59)^184^. To create the mutant co-fitness matrix, pairwise Pearson correlations for the chemical-genetic profiles of each of the 1909 mutants above were calculated and were hierarchically clustered using centroid clustering with Euclidean distance as the distance metric. The resulting heatmaps were visualized and adjusted in TreeView (version 1.2.0)^185^.

### Random Forest classifier and feature importance analysis

We designed a machine learning pipeline to infer mutants’ *in vivo* attenuation based on *in vitro* phenotypes (Fig. 4F). We defined the 157 *in vitro* relative fitness scores for each mutant as features. We framed the *in vivo* prediction as a binary classification problem, defining two classes that we refer to as “Attenuated” and “Not Attenuated.” Specifically, the “Attenuated” class corresponds to mutants with an *in vivo* relative fitness score less than or equal to −2, while “Not Attenuated” is defined for *in vivo* relative fitness scores strictly higher than 0. We excluded stressors with *in vivo* relative fitness scores in the interval (−2,0], as this range may contain false negative attenuated mutants that could complicate classification. Thus, our set of training data can be represented as {x_l_, y_l_ } where x_l_ and y_l_ are the set of 157 features and the binary target label for the mutant *i*, respectively, with y*_i_* = 0 if the target class is “Attenuated,” or y*_i_* = 1 if it is “Not Attenuated.”

We employed a Random Forest (RF) as a classification model. For training and evaluating the RF model, we performed a leave-one-out approach on the 10 non-overlapping subpools of mutants that were profiled independently. Specifically, we trained 10 models, each time exploiting 9 subpools as the training set and the remaining subpool as a test set to evaluate our classifier’s accuracy based on prediction of mutants in the unseen subpool as “Attenuated” or “Not Attenuated” (the target label). To further increase the generality assessment, for each leave-one-out experiment we computed our target labels from one random mouse, among the five available in each pool. This design choice is motivated by the necessity to evaluate the robustness of the classification algorithm with respect to all the available different mutants and assessing the generality of the proposed model.

The choice of an RF classifier was motivated by the possibility to measure feature importance. RF is designed as a parallel set of decision trees that are trained on random splits of data and features (i.e., a bootstrapped ensemble), and whose predictions are averaged together to obtain the final prediction. Mean Decrease in Impurity (MDI) is a measure used to provide a feature importance score in the RF classification procedure. Here “Impurity” refers to the extent to which the data at a single node in the decision tree tend to belong to the same class (High Purity/Low Impurity) or belong to many different training classes (Low Purity/High Impurity). This is related to the Gini Purity (or, in some implementations, Information Entropy), used to decide onto which features a specific node of the Decision Tree should make the split. In a single decision tree, the feature to be selected is the one that reduces impurity the most, until it ideally reaches 0 at leaf nodes. MDI is obtained by considering a specific feature x_k_ and summing the decrease in Impurity obtained by traversing the decision tree according to that feature. This is computed for all features and for all nodes. Then, it is aggregated across all the Decision Trees in the Random Forest to obtain the final MDI scores. Feature importances are normalized to sum up to one.

### Protein conservation and domain analysis

Ortholog groups and predicted domains for the *C. neoformans* proteome were accessed via FungiDB (release 66)^186^. Ortholog groups were searched in OrthoMCL (release 6.21)^12^ and were classified as being specific to pathogenic *Cryptococcus* spp. if that group was found only in the *C. neoformans*, *C. deneoformans*, *C. gattii*, *C. deuterogattii*, *C. bacillisporus*, or *C. tetragattii* genomes present in the database. Proteins found beyond these pathogenic species but restricted to tremellomycetes were classified as being tremellomycete-specific, and so on to categorize proteins as basidiomycete-specific, fungus-specific, eukaryote-specific, or conserved in bacteria and/or archaea. We used information from Interpro, PFam, PirSF, Prosite, Smart ID, Superfamily, and TigrFam databases (accessed via FungiDB) to determine whether proteins had at least one bioinformatically predicted protein domain.

### Yeast spot dilution assays

Overnight yeast cultures were harvested, washed once in water, resuspended in water, and adjusted to 3 x 10^6^ yeast/mL. Suspensions were 5-fold serially diluted five times to obtain six resuspensions ranging in concentration from 960 yeast/mL to 3 x 10^6^ yeast/mL. 3 μL of each dilution was spotted onto an agar plate and allowed to dry. Plates were incubated for 2-3 days before imaging.

### Growth for Rim101 western blots

To assess Rim101 processing in wild-type, *rra1Δ*, or *rra2Δ* backgrounds, overnight cultures grown in YPAD were inoculated into unbuffered YPAD or YPAD buffered to pH 8.0 with 100 mM HEPES-NaOH and were cultured for 4 hours before harvesting for protein extraction and western blotting as described below. To assess Rim101 processing in response to different Rra1 alleles, cultures were grown in unbuffered SC to OD_600_ ∼1.0, diluted in SC buffered to the desired pH with McIlvaine’s buffer, and cultured for 4 hours before harvesting for western blotting.

### Protein extraction

Logarithmically growing yeast (2 OD_600_’s) were pelleted, washed once with water, and stored at −70°C until later processing. Pellets were resuspended in 200 μL 10% TCA, incubated on ice for 10 min, and pelleted at 21,000 x*g* at 4°C for 5 min. The supernatant was discarded, and pellets were washed with cold acetone and centrifuged at 21,000 x*g* at 4°C for 5 min. Pellets were air dried and resuspended in 200 μL 2X NuPAGE LDS sample buffer (Invitrogen NP0007) supplemented with 50 mM Tris-HCl, pH 8.0 and 100 mM DTT. Samples were then bead-beat on an Omni Bead Ruptor Elite with 0.5 mm zirconia/silica beads at 6.0 m/s for 2 cycles of 90 s with a 90 s rest between, followed by boiling for 10 min.

### Western blotting

Samples were separated on 4-12% Bis-Tris gels (Genscript) in Tris-MOPS-SDS running buffer (Genscript M00138) and transferred to 0.45 μm nitrocellulose in Tris-Glycine transfer buffer (25 mM Tris, 250 mM glycine) with 20% methanol. Membranes were blocked in 5% milk in TBST (10 mM Tris-HCl, pH 7.4, 150 mM NaCl, 0.1% Tween-20) for 1 hr at room temperature or overnight at 4°C followed by incubation in primary antibody (1:3000 mouse-α-FLAG M2, Sigma F3165, or 1:1000 rabbit-α-Histone H3, Invitrogen PA5-16183) in blocking buffer for 1 hr at room temperature or overnight at 4°C. Membranes were washed three times in TBST and incubated in HRP-conjugated secondary antibody (1:8000 goat-α-mouse IgG, Invitrogen 31430, or 1:8000 goat-α-rabbit IgG, Invitrogen 65-6120) for 1 hr at room temperature followed by three washes in TBST. Membranes were rinsed in PBS and detected with SuperSignal West Pico PLUS (Thermo Scientific 34580) or SuperSignal West Femto Maximum Sensitivity (Thermo Scientific 34094) chemiluminescent substrate.

### Microscopy

For Rim101, Rim23, Rra1, and Rra2 imaging experiments, overnight YPAD cultures were diluted to OD_600_ 0.1 in SC buffered to pH 4.0 with McIlvaine’s buffer and were cultured 4-6 hours to OD_600_ ∼0.4-0.6. Cultures were then pelleted, resuspended in SC buffered to either pH 4.0 or pH 8.0 with McIlvaine’s buffer, and cultured for 30 minutes. Yeast were spotted onto 2% agarose pads (Lonza 50081) made in the same medium as the liquid culture (SC buffered to pH 4.0 or 8.0, as appropriate), allowed to dry, and covered with #1.5 coverslips prior to live imaging. For Cps1 imaging, overnight YPAD cultures were diluted to OD_600_ 0.1 in YNB, grown for 3-4 hours to OD_600_ ∼0.4-0.5, and spotted onto 2% agarose pads in YNB prior to mounting and live imaging.

Confocal images were captured on an inverted Nikon Ti microscope equipped with a CSU-22 spinning disk (Yokogawa) and an Evolve Delta EMCCD camera (Photometrics). Imaging was performed with a Plan Apo VC 100X/1.4 NA oil objective using a 488 nm solid-state laser for excitation and an ET525/50m emission filter for mNeonGreen. Brightness and contrast were adjusted in Fiji^187^. Image acquisition and processing parameters were constant within each experiment.

### Growth and harvesting for affinity purification-mass spectrometry

For Rra2, Mid1, Unc79, Unc80, Rad9, Njf1, and Cps1 AP-MS, strains were inoculated into 2 L YPAD at a low density (OD_600_ ∼0.001-0.002) and were grown shaking at 200 rpm overnight at 30°C. The next day, at OD_600_ ∼1.0, 1% additional glucose was added. For Mid1, Unc79, Unc80, and Cps1, cultures were then grown directly to OD_600_ ∼2.0 and harvested. For Rra2, the medium was buffered shortly after glucose addition with 100 mM (final concentration) HEPES-HCl, pH 4.0 or 100 mM HEPES-NaOH, pH 8.2. Cultures were then grown for 1 hr in the presence of buffer before harvest. For drug-treated Rad9 and Njf1 samples, 500 μ/mL Zeocin was added concurrently with glucose addition, and cells were grown for 2 additional hours before harvest. For Rgh1 AP-MS, overnight YPAD cultures were washed in DMEM, pH 7.4, diluted to an OD_600_ of 0.2 in DMEM, pH 7.4, and cultured shaking at 200 rpm at 37°C without CO_2_ until harvest at OD_600_ ∼1.0. For all samples, cells were harvested by centrifugation at 5000-6200 xg at 4°C for 5-10 min in a JLA-8.1000 rotor, washed with 300 mL cold water, and pelleted at 9000-10,500 xg for 5-10 min at 4°C. Rgh1 samples were washed with 300 mL of a modified immunoprecipitation (IP) wash buffer (25 mM HEPES-KOH, pH 7.9, 300 mM KCl) instead of water, as yeast grown in DMEM do not readily pellet after water washes. Samples were resuspended in the appropriate IP buffer (see below) and were stored at −70°C as “popcorn” at described below.

### Membrane pellet affinity purification-mass spectrometry

For IPs performed from membrane pellets (Rra2, Mid1, Unc79, Unc80, and Cps1), pellets were resuspended in 15-20 mL Membrane IP buffer (50 mM HEPES-KOH, pH 7.9, 150 mM KOAc, 2 mM MgOAc, 1 mM CaCl_2_, 200 mM sorbitol, 2 mg/mL iodoacetamide, and protease inhibitors [Pierce A32965 or Sigma 11836153001]). Resuspended pellets were dripped slowly into liquid nitrogen to form “popcorn” and were stored at −70°C until further processing. Cells were lysed by cryogrinding in a SPEX Sample Prep 6870 for 9-12 cycles of 3 min grinding at 15 cps followed by 2 min cooling between cycles. Samples were stored at −70°C until further processing.

Grindates were thawed and treated with 2500 U Benzonase (Sigma E1014) or Benz-Neburase (GenScript Z03695) for 30-90 min at room temperature. Samples were pelleted at 16,000 xg at 4°C for 15 min in a JA-17 rotor and the supernatant was discarded. The cell pellet was resuspended in 10 mL Membrane Solubilization buffer (50 mM HEPES-KOH, pH 7.9, 250 mM KOAc, 2 mM MgOAc, 1 mM CaCl_2_, 200 mM sorbitol, 15% glycerol, 1% (w/v) GDN, 0.1% CHS, 2 mg/mL iodoacetamide and protease inhibitors), incubated for 2 h at 4°C with agitation, and centrifuged at 20,000 xg for 45 min at 4°C. The solubilized supernatant was collected for anti-FLAG IP. Meanwhile, 400 μL anti-FLAG M2 Magnetic beads (Sigma M8823) were prepared by 3 washes (10 min with rocking at 4°C) with Membrane IP Wash buffer (50 mM HEPES-KOH, pH 7.9, 250 mM KOAc, 2 mM MgOAc, 1 mM CaCl_2_, 200 mM sorbitol, 15% glycerol, 0.1% GDN, 0.01% CHS and protease inhibitors). The solubilized supernatant was added to beads and incubated overnight at 4°C with rocking. The flow-through was discarded, and non-specific material removed by 3 washes with 15 mL Membrane IP Wash buffer followed by 3 washes with Low-detergent Membrane IP Wash buffer (50 mM HEPES-KOH, pH 7.9, 250 mM KOAc, 2 mM MgOAc, 1 mM CaCl_2_, 200 mM sorbitol, 15% glycerol, 0.01% GDN). Samples were eluted by 3-4 incubations in 200-500 μg/mL 3X FLAG peptide (Sigma F4799) in Low-detergent Membrane IP Wash buffer in 600-900 μL final volume. For Cps1 samples, samples were washed and eluted in Detergent-free Membrane IP Wash buffer (lacking GDN or CHS) instead of Low-detergent Membrane IP Wash buffer. Eluates were pooled, TCA was added to 13% final concentration to precipitate proteins, and samples were incubated on ice overnight at 4°C. Proteins were pelleted at 21,000 xg for 20 min at 4°C, washed 2-3 times with ice-cold acetone, and stored at - 70°C until further processing.

TCA pellets were prepared for MS using the PreOmics iST Sample preparation kit according to the manufacturer instructions. Briefly, samples were reduced and alkylated by incubation in Lysis buffer for 10 min at 95°C with shaking followed by Trypsin/LysC digestion for 3 hr at 37°C with shaking. The reaction was stopped with Stop buffer, and samples were loaded onto a cartridge and spun through (2250 xg for 3 min). Cartridges were washed sequentially with Wash X, Wash 1, and Wash 2, followed by elution and vacuum evaporation until dry.

### Non-membrane pellet affinity purification mass spectrometry

For non-membrane pellet AP-MS samples (Rad9, Njf1, and Rgh1), harvested cell pellets were resuspended in 15-20 mL Standard IP buffer (25 mM HEPES-KOH, pH 7.9, 300 mM KCl, 0.1 mM EDTA, 0.5 mM EGTA, 2 mM MgCl_2_, 20% glycerol, 0.1% Tween-20, 1 mM DTT) with protease inhibitors. DTT was omitted from all buffers for Rgh1 samples, except for elution buffer, to minimize disruption of this protein’s predicted disulfide bonds. “Popcorn”-making, cryogrinding, and Benzonase/Benz-Neburase treatment was performed as described above. Whole cell lysates were clarified by centrifugation at 40,000 xg at 4°C for 15 min in a JA-17 rotor, and the supernatant was collected for anti-FLAG IP. Meanwhile, 400 μL anti-FLAG M2 Magnetic beads were prepared by 3 washes (10 min with rocking at 4°C) with Standard IP buffer. The clarified lysate was added to beads and incubated overnight at 4°C with rocking. The flow-through was discarded, and non-specific material removed by 3 washes with 15 mL Standard IP buffer followed by 3 washes with Detergent-free Standard IP buffer (Standard IP buffer without Tween-20). Samples were eluted by 3 x 300 μL incubations in 200-500 μg/mL 3X FLAG peptide in Detergent-free Standard IP buffer. Eluates were TCA precipitated and reduced, alkylated, and digested as described above, except that Wash X was omitted.

### Hyaluronic acid (HA) ELISA

HA production was measured using the HA-ELISA kit (Corgenix 029-001). Yeast were cultured in YPAD to OD_600_ ∼0.4-0.6, washed with sterile PBS, and adjusted to 5 x 10^8^ CFU/mL in PBS. 10 μL (5 x 10^6^ CFU) were diluted in 100 μL Reaction Buffer and incubated in HA-binding protein (HABP)-coated microwells overnight at room temperature with shaking. The next day, wells were washed with Wash Solution to remove yeast, HRP-conjugated HABP Solution was added, and wells were incubated for 30 min at room temperature. Wells were washed, and One-component Substrate Solution (containing 3,3’,5,5’-tetramethylbenzidine and H_2_O_2_) was added and incubated for 30 min for color development. The reaction was stopped with Stopping Solution (0.36 N sulfuric acid), and intensity was measured at 450 nm with a 650 nm reference. Final HA concentrations were calculated by comparing with an HA standard curve and reference solutions supplied with the kit. Statistical differences were analyzed in GraphPad Prism (version 10.2.3).

### *RGH1* qRT-PCR

Overnight YPAD cultures of wild-type yeast were washed in PBS and inoculated into YPAD, YNB, or DMEM (containing sodium bicarbonate). For YPAD and YNB, yeast were inoculated at an initial OD_600_ of 0.1 and were cultured at 30°C on a roller drum until harvest at OD_600_ ∼1.0. For DMEM, yeast were inoculated at an initial OD_600_ of 2.5 in 5 mL and were cultured static for 6 hr in 6-well tissue culture plates at 37°C/5% CO_2_. Samples for RNA isolation were harvested by centrifugation, frozen in liquid nitrogen, and stored at −70°C until further processing.

To isolate RNA, 500 μL TRIzol (Invitrogen 15596018) and 100 μL 0.5 mm zirconia/silica beads were added to frozen pellets. Samples were bead-beaten on an Omni Bead Ruptor 12 (Omni International 19-050) on “high” for 90 s and were incubated on ice for 5 min. 100 μL chloroform was added, and the sample was mixed by inversion and incubated at room temperature for 10 minutes. Samples were centrifuged for 15 min at 21,000 xg at 4°C, and the aqueous phase was collected. An equal volume of ethanol was added, and RNA was isolated with an RNA Clean & Concentrator-5 kit (Zymo R1013). Briefly, the sample was passed over a Zymo-Spin IC column and was treated on-column with 6 U TURBO DNase (ThermoFisher AM2238) at 37°C for 30 min. RNA was washed and eluted in 20 μL nuclease-free water according to manufacturer’s instructions. cDNA was synthesized from 1 μg total RNA with SuperScript III reverse transcriptase (Invitrogen 18080-044) and random hexamers according to manufacturer’s instructions. cDNA samples were diluted 1:5 with water, and 2.5 μL were used as templates for qPCR with PowerUp SYBR Green master mix (Life Technologies A25742) using 500 nM each forward and reverse primers and a 10 μL reaction volume. Samples were analyzed on a CFX96 Real-Time PCR system (Bio-Rad) with initial denaturation of 50°C for 2 min and 95°C for 2 min followed by 40 cycles of 95°C for 15 s, 55°C for 15 s, and 72°C for 1 min. Each biological replicate was analyzed in technical triplicate, and relative expression differences were calculated using the ΔΔC_t_ method with actin as a reference.

### Secretome profiling

For YNB culture supernatant and cell pellet proteomics, overnight YPAD cultures were inoculated to an OD_600_ of 0.001 in 200 mL of YNB and were grown for 24 hours shaking at 200 rpm at 30°C. For DMEM samples, overnight YPAD cultures were inoculated to an initial OD_600_ of 0.2 in 5 x 50 mL cultures (250 mL total per biological replicate) in 15 cm tissue culture dishes. Cultures were incubated static for 24 hours at 37°C/5% CO_2_.

Harvested supernatants were passed through a 0.22 μm filter followed by concentration on a 3 kDa molecular weight cutoff Centricon Plus-70 filter (Millipore Sigma UFC700308). Samples were then precipitated with TCA (10% final concentration) and were washed twice with 10% TCA and twice with cold acetone. Pellets were stored at −70°C until further processing with the PreOmics iST Sample preparation kit as described above.

Cell pellets (0.25 g for YNB, 1.0 g for DMEM) were processed as previously described^188^ with modifications. Briefly, pellets were resuspended in 5 mL 100 mM Tris-HCl, pH 8.5 containing protease inhibitors and were lysed using a probe sonicator at 30% power. We used 2 cycles of 30 s on/30 s off for YNB cell pellets and 5 cycles for DMEM cell pellets. Lysates were pelleted for 30 min at 4°C at 20,000 xg in a JA-17 rotor, and supernatants were harvested. SDS (2% final concentration) and DTT (10 mM final concentration) were added, and samples were incubated at 95°C for 10 min. Samples were cooled to room temperature, iodoacetamide (0.55 mM final concentration) was added, and samples were incubated in the dark for 10 min. Samples were then TCA precipitated and stored at −70°C until further processing for mass spectrometry with the PreOmics iST Sample preparation kit as described above.

### Mass spectrometry data acquisition and search parameters

A nanoElute was attached in line to a timsTOF Pro equipped with a CaptiveSpray Source (Bruker, Hamburg, Germany). Chromatography was conducted at 40°C through a 25 cm reversed-phase C18 column (PepSep, 1893476) at a constant flowrate of 0.5 μL/min. Mobile phase A was 98/2/0.1% water/MeCN/formic acid (v/v/v, Thermo Fisher, LS118, LS120) and phase B was MeCN with 0.1% formic acid (v/v). During a 108 min method, peptides were separated by a 3-step linear gradient (5% to 30% B over 90 min, 30% to 35% B over 10 min, 35% to 95% B over 4 min) followed by a 4 min isocratic flush at 95% before washing and a return to low organic conditions. Experiments were run as data-dependent acquisitions with ion mobility activated in PASEF mode. MS and MS/MS spectra were collected with m/z 100 to 1700 and ions with z = +1 were excluded.

Raw data files were searched using PEAKS Online Xpro 1.7.2022-08-03_160501 (Bioinformatics Solutions Inc., Waterloo, Ontario, Canada). The precursor mass error tolerance and fragment mass error tolerance were set to 20 ppm and 0.03 respectively. The trypsin+LysC digest mode was set to semi-specific and missed cleavages was set to 3. We searched the data using the FungiDB database. Carbamidomethylation was selected as a fixed modification. Oxidation (M), Deamidation (NQ), and Acetylation (N-term) were selected as variable modifications.

### Affinity purification-mass spectrometry data analysis

Datasets were filtered to exclude contaminants and proteins for which MS1 area values were not obtained for any of the samples within a given comparison. A pseudovalue of 1 was added to all protein-level MS1 area values to enable downstream log_2_ transformation. MS1 areas were normalized to the sum of total MS1 area within a sample, and values for biological replicates were averaged. Protein enrichments reported on graphs are log_2_-transformed ratios (tagged/untagged) average MS1 areas.

### Supernatant proteomics data analysis

Samples from cells grown in DMEM were labeled “DP” for cell pellets and “DS” for cell supernatants. Those from cells grown in YNB were labeled “YP” for cell pellets and “YS” for supernatants. The MS data were filtered for proteins for which at least 10 spectra were obtained across all experiments. Proteins without MS1 area data for any of the supernatants were filtered out. For the samples grown in DMEM, replicate 4 (labeled “DS_4”) for the supernatant was dropped from analysis due to low signal. The same was done for replicate 4 of the supernatants for samples grown in YNB (labeled “YS_4”). For the remaining data, the proteins were sorted based on the ratio of the sum of the MS1 area values for the supernatants divided by those of the pellets. For the moving χ^2^ analysis, for each possible cutoff, the sum of the supernatant MS1 areas above and below the cutoff was computed, and the same was done for the sum of the MS1 area values for the pellets. On that 2 x 2 table, the χ^2^ statistic was calculated using the chi2_contingency function in the stats module from SciPy. This was done for all possible cutoffs. The cutoff producing the maximum χ^2^ value was selected to define proteins as secreted.

### CRISPR suppressor screen growth

To identify loss-of-function suppressors of the *rgh1Δ* phenotype, we created a genome-scale insertional mutagenesis library in a Cas9-expressing, *rgh1Δ* strain. To do this, we utilized a plasmid library containing sgRNAs targeting 4977 *C. neoformans* genes with 4 guides per gene (M.Y.H. and H.D.M., unpublished data). These sgRNAs were cloned adjacent to a NAT resistance cassette and a UMI, enabling PCR amplification of fused NAT-sgRNA-UMI amplicons. We electroporated these amplicon pools in duplicate into Cas9-expressing, *rgh1Δ* yeast and cultured in YPAD containing 75 μg/mL nourseothricin to select for insertional mutants into the target loci. We obtained a library of approximately 2 x 10^5^ transformants representing ∼10X coverage per sgRNA. To identify suppressors, we inoculated 2 x 10^7^ CFUs (∼100X library coverage) from frozen library glycerol stocks of our duplicate libraries into DMEM, pH 7.4, and cultured shaking at 200 rpm at 37°C/5% CO_2_ for 4 days. Cultures were pelleted, inoculated into 100 mL YPAD cultures at an initial OD_600_ of 0.1, and outrgrown in YPAD for harvesting and gDNA isolation as described above.

### CRISPR suppressor screen sequencing library preparation

Concentration of gDNA samples was measured using a Qubit dsDNA BR Assay Kit. To prepare sequencing libraries, sgRNA and UMI sequences were PCR amplified from gDNA samples using custom primers carrying Illumina adapter and index sequences on overhangs. Briefly, 15 cycles of PCR were run using NEBNext Ultra II Q5 Master Mix. To preserve library diversity, gDNA quantity equivalent to approximately 4 x 10^7^ genomes was used as template for each sample. PCR products were cleaned using a Monarch PCR and DNA cleanup kit (NEB T1030) following manufacturer’s instructions. Library size was analyzed using TapeStation High Sensitivity D1000 Screentape. The library was sequenced on a NextSeq 500 using a 75 cycle Nextseq 500/550 High Output v2.5 kit (Illumina 20024906) with a custom sequencing primer (P02) spiked in, as well as a 5% PhiX sample spike in.

### CRISPR suppressor screen data analysis pipeline

Reads were basecalled using bcl2fastq (version 2.17.1.14) with options –minimum-trimmed-read-length 10 –mask-short-adapter-reads 0 –barcode-mismatches 1 –no-lane-splitting. Excess cycles and low-quality reads were removed using Cutadapt (version 4.2)^178^ with options -q 25 -m 20:10 –max-n 0 -a CTAGATGCGAGATAT. sgRNA sequences were directly aligned to the expected library sequences using custom Python (version 3.10.8) scripts while tracking counts associated with each encountered UMI sequence for each sgRNA sequence. We reasoned that low counts associated with rare UMIs for each sgRNA were likely to arise from PCR contamination. We employed the following strategy to remove these reads. We first collapsed neighboring UMI sequences using the UMIClusterer function with cluster_method = ‘directional’ within UMItools (version 1.1.2)^181^ the Python API. We then applied a read count cutoff determined by the maximum of either 2 or log_2_(reads + pseudocount) - 2. Any reads below the cutoff were discarded. Reads were then analyzed by Mageck (version 0.5.9.5)^15^ using mageck mle with options –norm-method total –no-permutation-by-group.

## Data availability

All raw mass spectrometry data is available for download on MassIVE, a full member of the ProteomeExchange, under the identifier: MSV000096014. Raw sequencing data are available from the NCBI Sequence Read Archive under BioProject IDs PRJNA1173758 and PRJNA1175672.

## Acknowledgments

We thank Sandy Johnson for critical reading of the manuscript. We thank Aashish Manglik, Robert Stroud, and Dave Toczyski for helpful discussions. This work was supported by R01AI00272 to H.D.M. M.J.B. was supported by the UCSF Multidisciplinary Training Program in Lung Disease (T32HL007185) and postdoctoral fellowships from the NIH (F32AI152270) and the American Heart Association (24POST1194861). M.Y.H. was supported by the UCSF Microbial Pathogenesis and Host Defense training grant (T32AI060537). Sequencing was performed at the UCSF Center for Advanced Technology (CAT), supported by UCSF PBBR, RRP IMIA, and NIH S10OD028511 grants, and at the Gladstone Genomics Core. Microscopy was performed at the UCSF Center for Advanced Light Microscopy (CALM).

## Author contributions

I.C., A.I.G., C.M.H., Y.M., S.P., M.C.R, and Y.X. constructed and validated the *C. neoformans* knockout collection. M.J.B. developed KO-seq, generated the phenotypic atlas of *C. neoformans in vitro*, performed the pooled infection screens, analyzed the data, and performed the experiments shown in Figs. 1-4 and 6, unless otherwise noted. S.B. performed the experiments shown in Fig. 5 and the affinity purification experiments shown in Fig. 2I. M.B.J. performed the experiments shown in Fig. 7 and the affinity purification experiments shown in Fig. 3L. A.L.W. performed the experiments in Fig. 2D and 2L, generated strains, and assisted with mouse experiments. M.Y.H. shared unpublished data and resources and performed the sequencing and data analysis for the CRISPR/Cas9 suppressor screen in Fig. 6I. S.L. assisted with the mouse experiment in Fig. 6D-F. M.C. and V.P.P. developed the random forest classifier. A.C. and A.L. performed mass spectrometry under the supervision of B.W.Z. T.D.G. performed and provided guidance on AlphaFold2 modeling. K.J.S. performed AlphaFold2 modeling and provided guidance on membrane protein purification experiments. M.J.B and H.D.M. wrote the paper with input from other authors. H.D.M. oversaw the work.

---

**Table S1. *C. neoformans* deletion library, related to Figure 1**.

**Table S2. *In vitro* growth conditions, related to Figure 1**.

**Table S3. Mutant phenotype data, related to** Figure 1.

**Table S4. Mutant cofitness matrix clusters, related to** Figure 1.

**Table S5. Affinity purification-mass spectrometry data, related to** Figures 2**, 3, 5, and 6.**

**Table S6. *In vivo* mutant fitness data, related to Figure 4**.

**Table S7. CRISPR/Cas9 suppressor screen data, related to** Figure 6.

**Table S8. *C. neoformans* secretome data, related to** **Figure 7**.

**Table S9. Primers, plasmids, and strains used in this study, related to** Figures 1-7.

**Figure S1.**
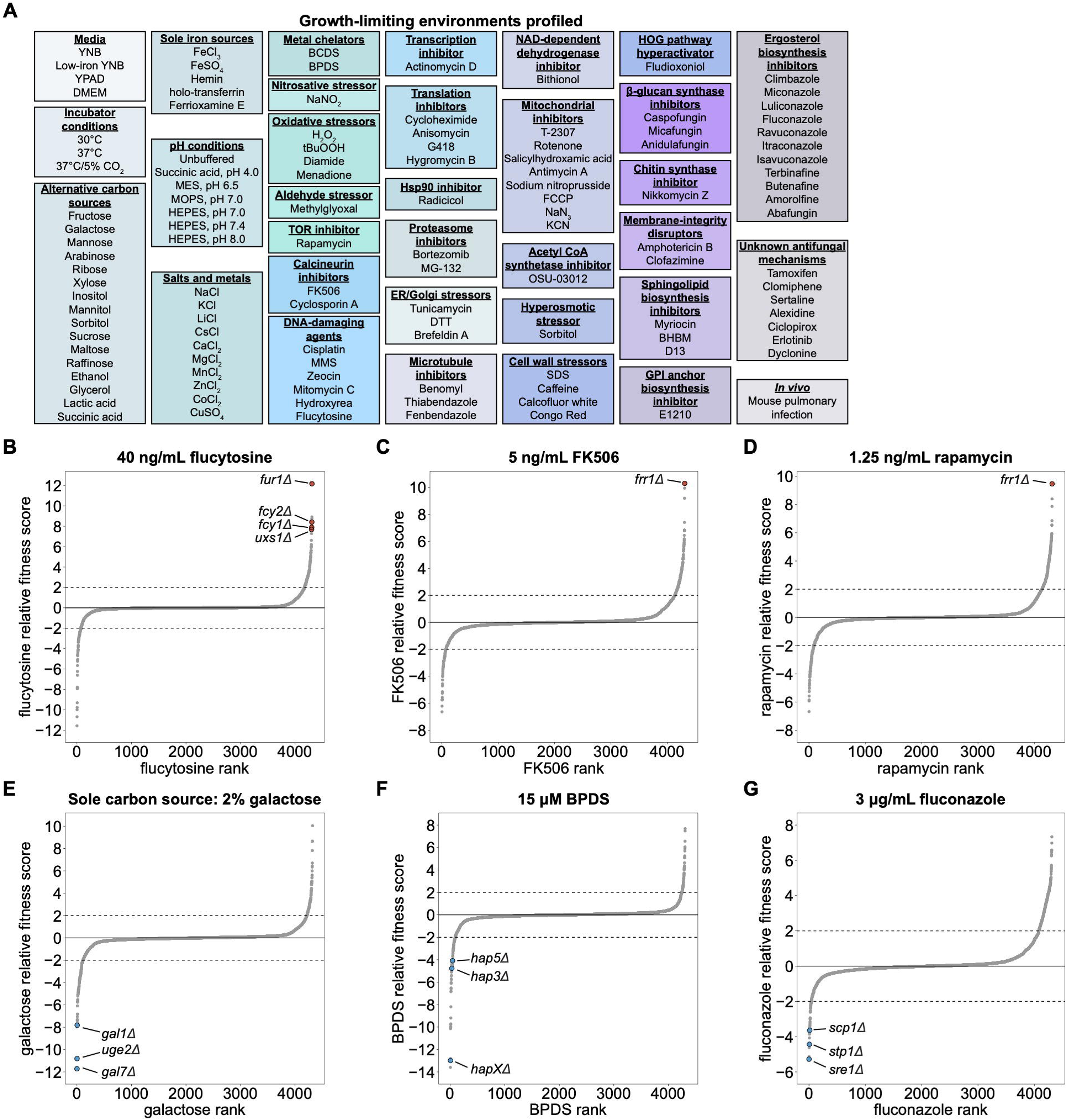
Pooled fitness profiling identifies determinants of stress sensitivity and tolerance, related to Figure 1. (A) Overview of growth environments profiled. (B-G) Rank-ordered plots of relative mutant fitness in example growth environments. Dashed lines indicate relative fitness scores of −2.0 and 2.0 that were used to define phenotypes. Mutants expected to display differential fitness in each condition based on prior studies are highlighted.

**Figure S2.**
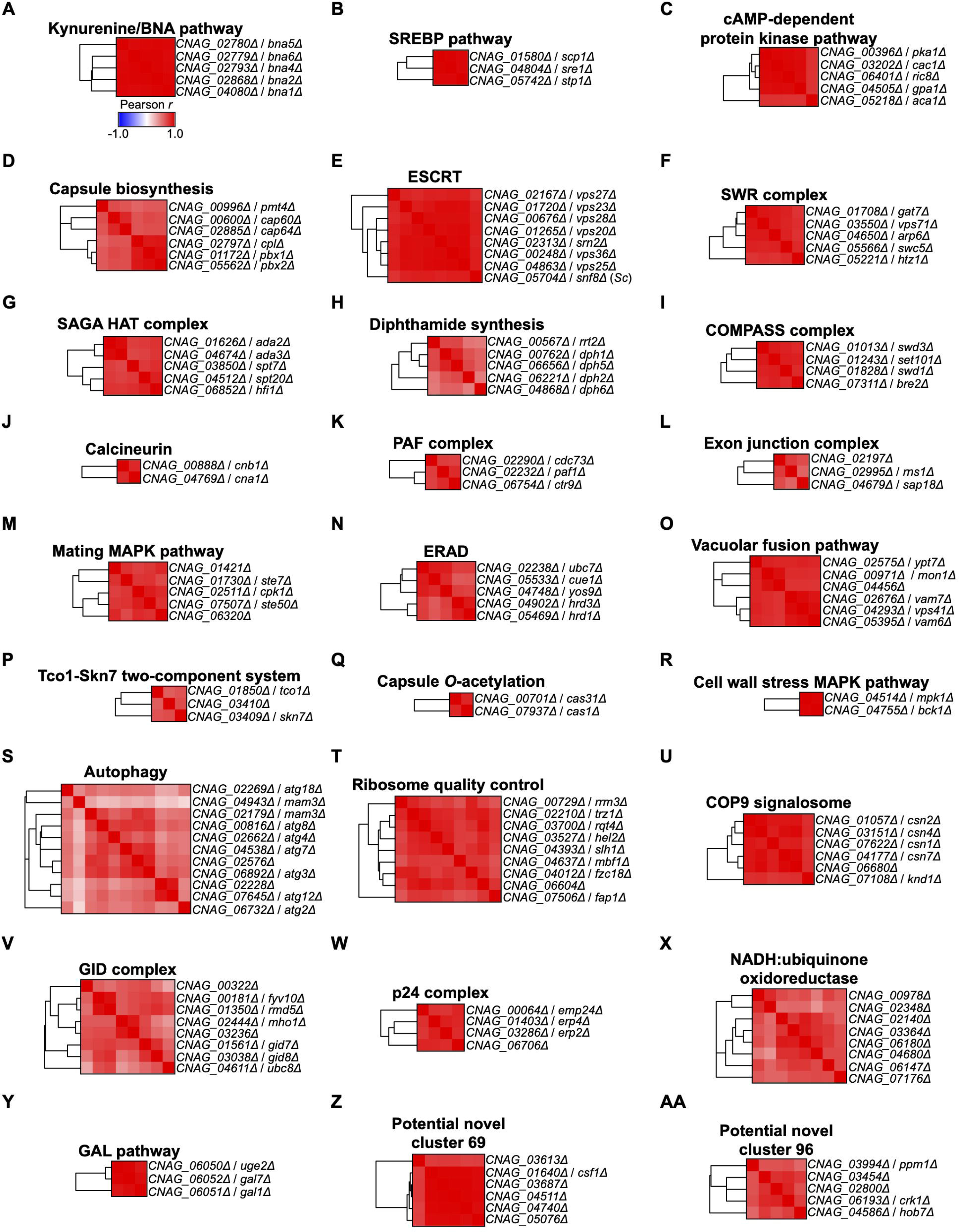
Mutant fitness profiling identifies functional genetic modules, related to Figure 1. (A-Y) Examples clusters of known function from Figure 1E. (Z, AA) Potentially novel clusters from Figure 1E. In cases where genes were not formally named in *C. neoformans*, gene names are derived from *S. cerevisiae* or *S. pombe* orthologs. Scale in (A) applies to all panels.

**Figure S3.**
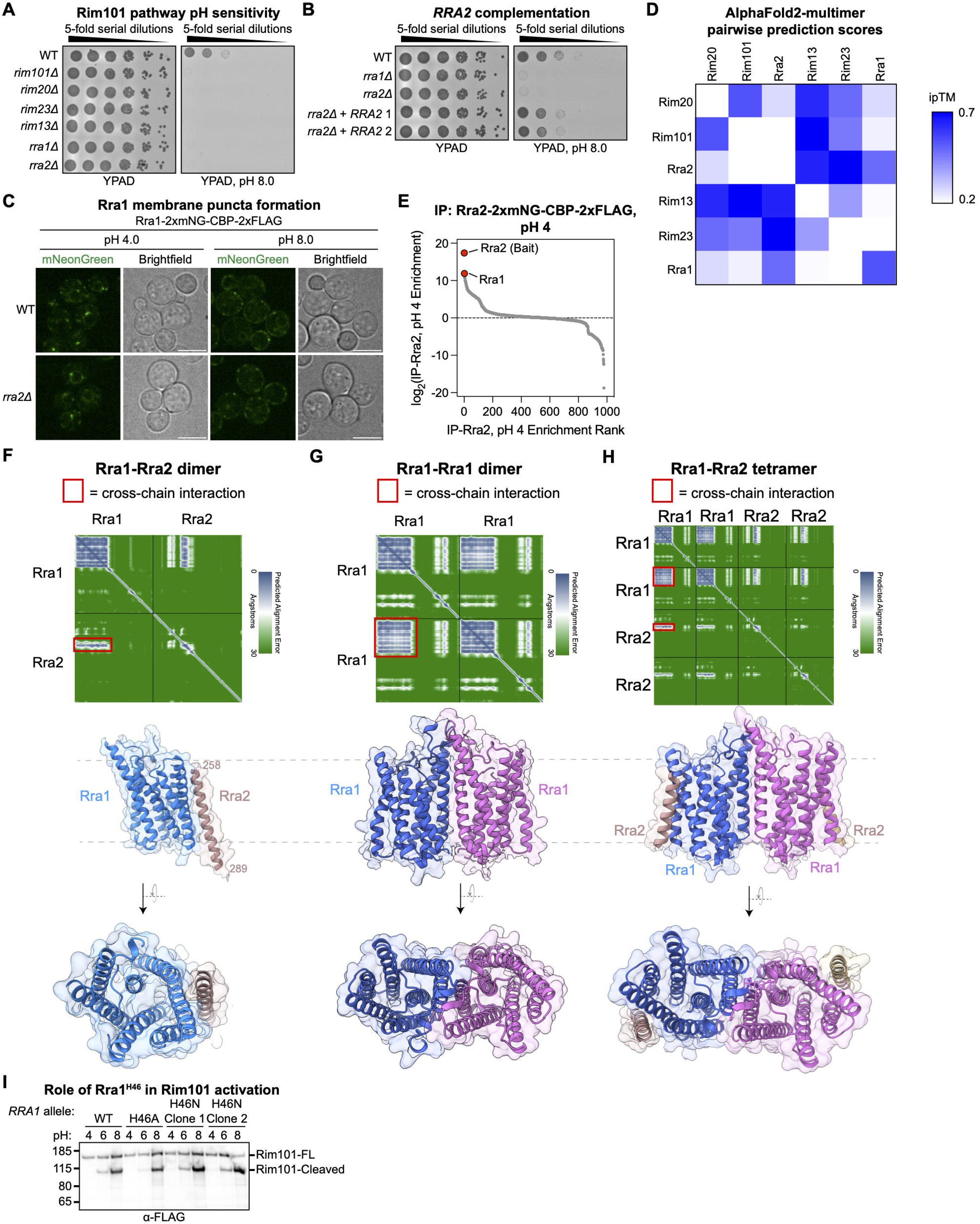
Additional data supporting a role for Rra2 in a pH-sensing complex, related to Figure 2. (A) Spot dilution assay of Rim101 pathway mutants in YPAD or YPAD, pH 8.0. Images were taken after 3 days of growth. (B) Spot dilution assays assessing complementation of the *rra2Δ* mutant by expression of a wild-type allele from the SH1 locus. Images were taken after 3 days of growth. (C) Live confocal imaging of endogenously tagged Rra1-2xmNG-CBP-2xFLAG in SC medium buffered to pH 4.0 or 8.0 with McIlvaine’s buffer. (D) Heatmap showing interface predicted template modeling (ipTM) scores for all pairwise AlphaFold2-multimer predictions of Rim101 pathway members. (E) Rra2 AP-MS at pH 4. YPAD cultures of yeast expressing endogenously tagged Rra2-2xmNG-CBP-2xFLAG were shifted to pH 4 for 1 hour, and anti-FLAG AP-MS was performed on membrane extracts. Data represent the protein-level ratio of normalized MS1 area from tagged versus untagged strains grown and processed in parallel. Averaged from 2 biological replicates. (F-H) AlphaFold2-multimer models and predicted alignment error plots for an Rra1-Rra2 dimer (F), an Rra1 homodimer (G), and an Rra1-Rra2 tetramer (H). (I) Independent biological replicate of experiment shown in Figure 2L assaying proteolytic processing of endogenously tagged 2xFLAG-mNG-CBP-Rim101 in response to *RRA1* alleles encoding His46 mutants.

**Figure S4.**
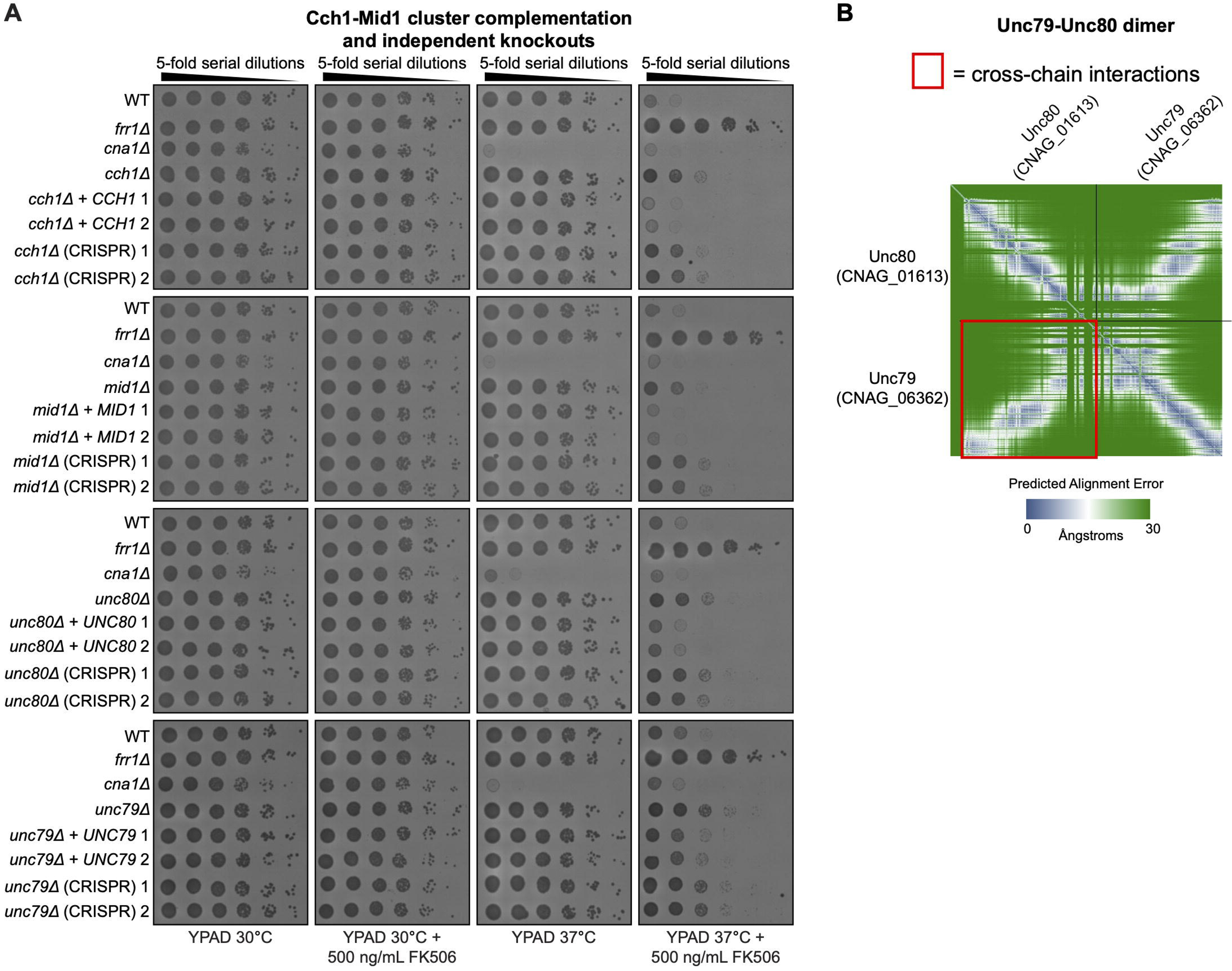
Additional data supporting roles for Unc79 and Unc80 in the Cch1-Mid1 pathway, related to Figure 3. (A) Spot dilution assays assessing FK506 sensitivity at 30°C and 37°C for Cch1-Mid1 cluster complement and independent CRISPR deletion strains. Images were taken after 2 days of growth. (B) Predicted alignment error plot for AlphaFold2-multimer Unc79-Unc80 dimer shown in Figure 3E.

**Figure S5.**
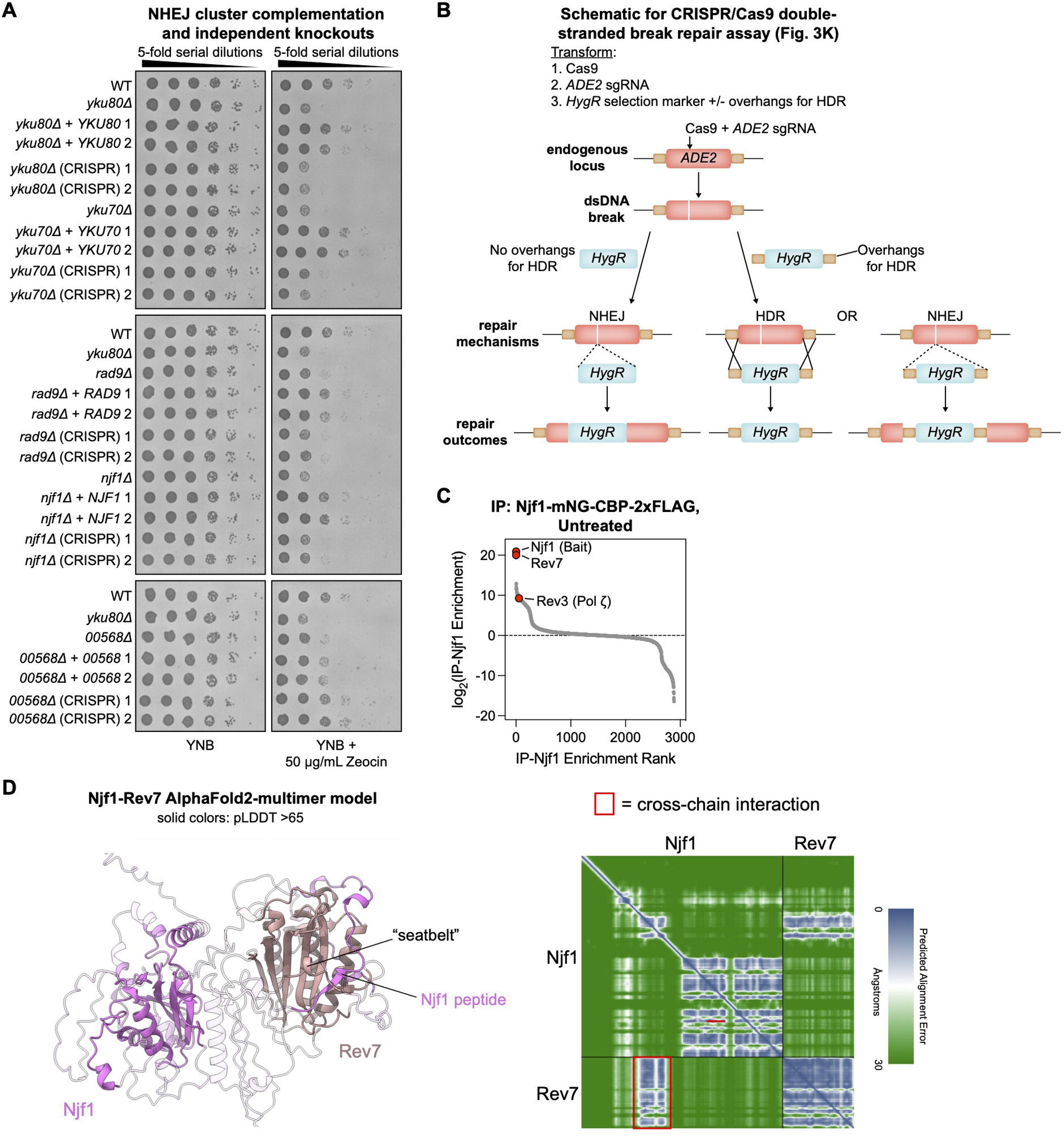
Additional data supporting roles for Rad9 and Njf1 in NHEJ, related to Figure 3. (A) Spot dilution assays assessing Zeocin sensitivity of knockout strains from the NHEJ cluster complemented with the wild-type gene and independent CRISPR-generated deletion strains. Images were taken after 2 days of growth. (B) Diagram of CRISPR/Cas9-induced double-stranded break repair assay shown in Figure 3K. (C) Njf1 AP-MS (untreated). Anti-FLAG AP-MS was performed on clarified cell lysates from a strain expressing endogenously tagged Njf1-mNG-CBP-2xFLAG. Data represent the protein-level ratio of normalized MS1 area from tagged versus untagged strains grown and processed in parallel. From 1 biological replicate. (D) AlphaFold2-multimer model and predicted alignment error for Njf1-Rev7 dimer. Confidently predicted protein segments are displayed in non-faded cartoon form.

**Figure S6.**
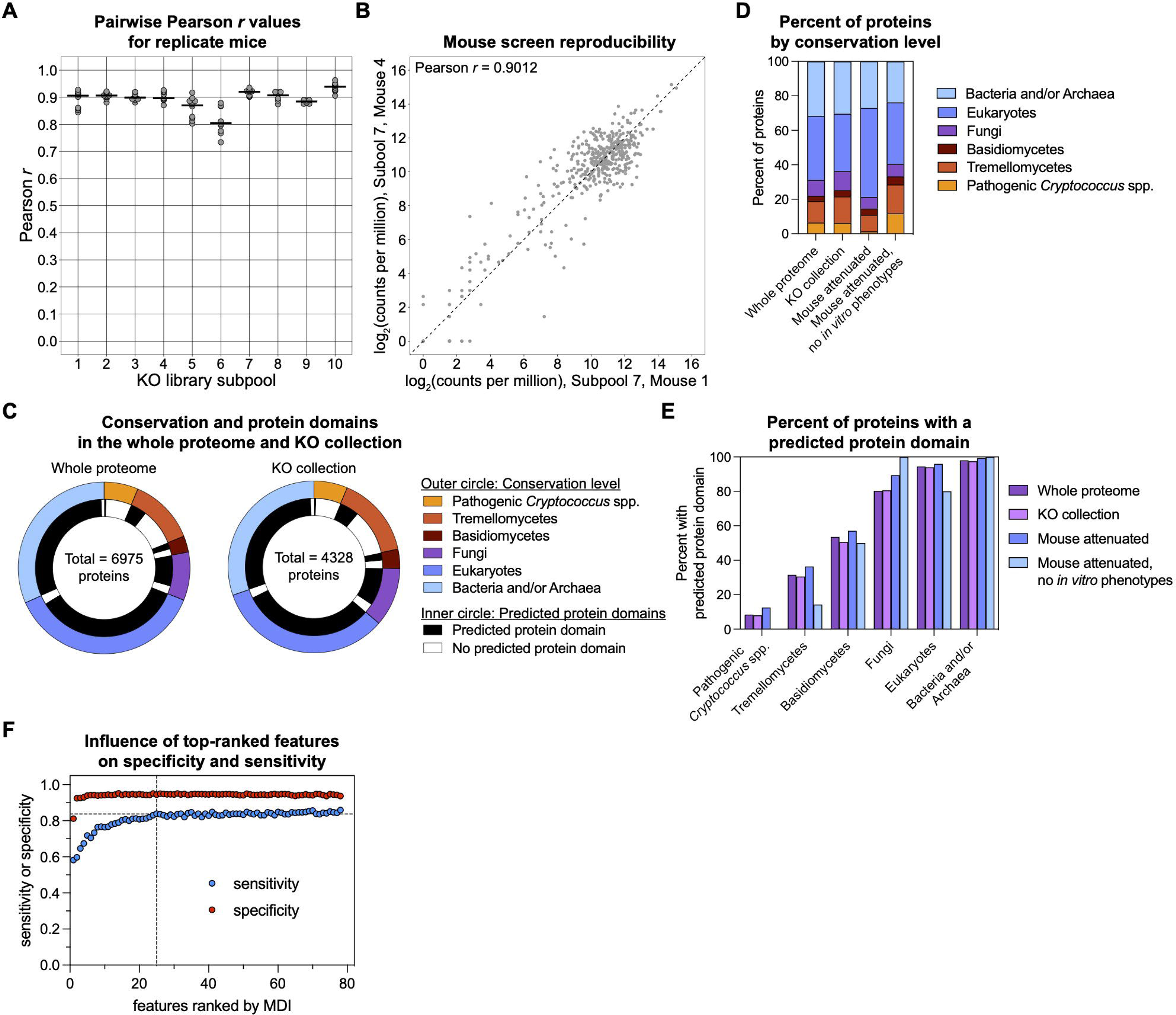
Additional data on infectivity-promoting genes, related to Figure 4. (A) Pearson correlation coefficients of mutant abundance in pairwise comparisons of replicate mice for the 10 library subpools screened. (B) Example comparison of mutant abundances in Subpool 7, Mouse 1, and Subpool 7, Mouse 4. These mice are the pair with the median Pearson *r* value across all comparisons in (A). (C) Conservation levels and predicted domains in proteins encoded by the full *C. neoformans* genome (left) and genes deleted in the knockout collection (right). Outer circle represents the most specific conservation level for a protein based on OrthoMCL orthology groups. Inner circle indicates the presence or absence of bioinformatically predicted protein domains. (D) Comparison of protein conservation levels between categories of interest from Fig. 4E and Fig. S6C. (E) Comparison of proteins with predicted domains by conservation group. (F) Impact of *in vitro* feature selection on sensitivity and specificity of *in vivo* phenotype prediction. Features were rank ordered by their importance (MDI), and their influence on sensitivity and specificity was plotted. Horizontal dashed line represents the average sensitivity of 0.837 achieved across all models. Vertical dashed line indicates the top 25 features that achieve this sensitivity.

**Figure S7.**
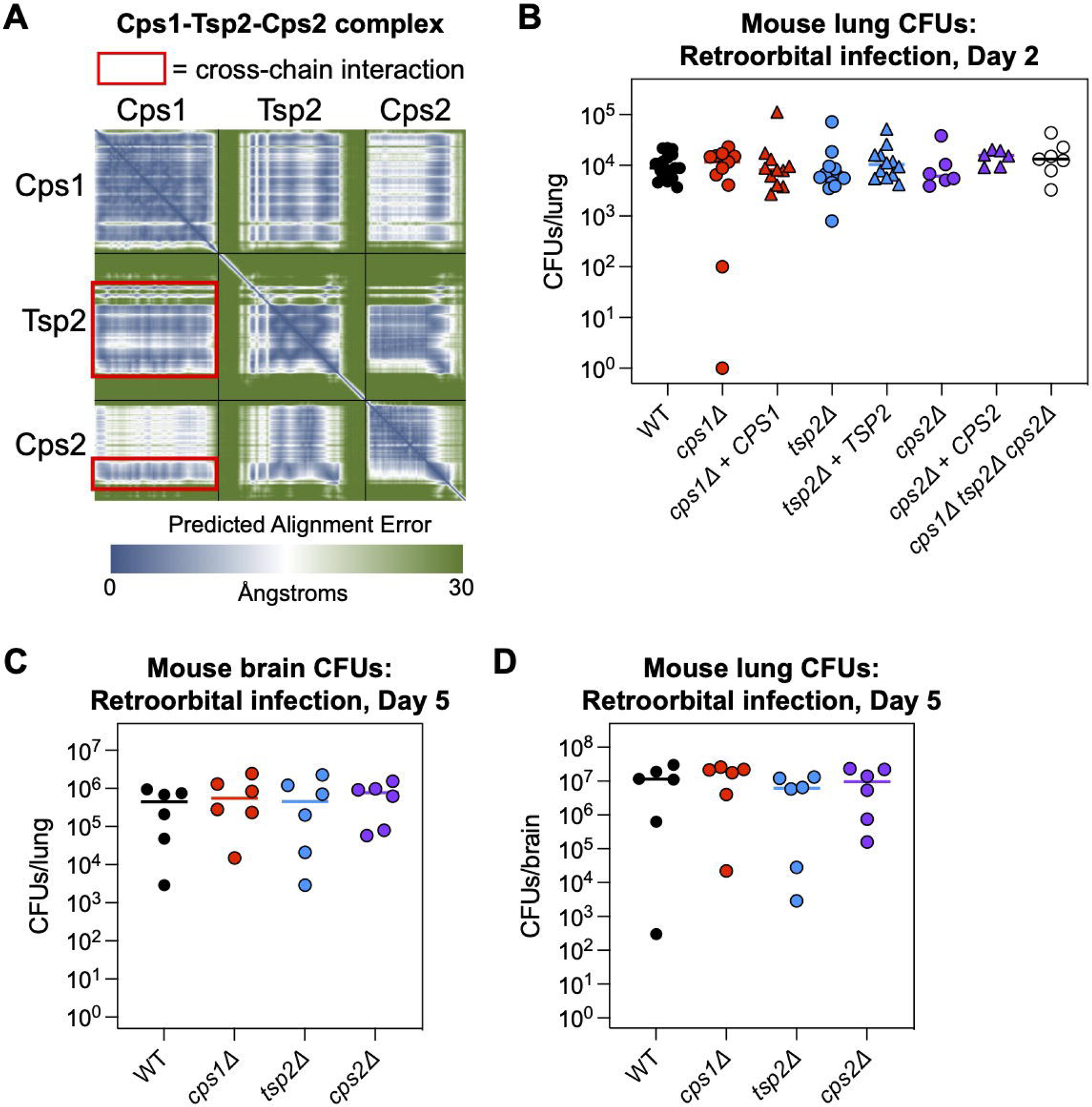
Additional data supporting a role for a tetraspanin-containing hyaluronic acid synthase complex in neurovirulence, related to Figure 5. (A) Predicted alignment error plot of the Cps1-Tsp2-Cps2 AlphaFold2-multimer model in Figure 5D. (B) Total lung CFUs 2 days post infection from mice infected intravenously (retroorbital inoculation) with 5 x 10^4^ CFUs of the indicated strains. Bars represent median values. WT, *n* = 17 mice pooled from 4 independent experiments with 3-6 mice per experiment; *cps1Δ* and *cps1Δ* + *CPS1*, *n* = 11 mice per group pooled from 3 independent experiments with 3-6 mice per experiment; *tsp2Δ* and *tsp2Δ* + *TSP2*, *n* = 12 mice per group pooled from 3 independent experiments with 3-6 mice per experiment; *cps2Δ*, *cps2Δ* + *CPS2*, and *cps1Δ tsp2Δ cps2Δ*, *n* = 6 mice per group pooled from 2 independent experiments with 3 mice per experiment. (C, D) Total brain (C) and lung (D) CFUs 5 days post infection from mice infected intravenously (retroorbital inoculation) with 5 x 10^4^ CFUs of the indicated strains. Bars represent median values, *n* = 6 mice per group from 1 experiment. Statistical analyses in (B-D) were performed with the Kruskal-Wallis test with Dunn’s multiple comparisons test. No significant differences were detected.

**Figure S8.**
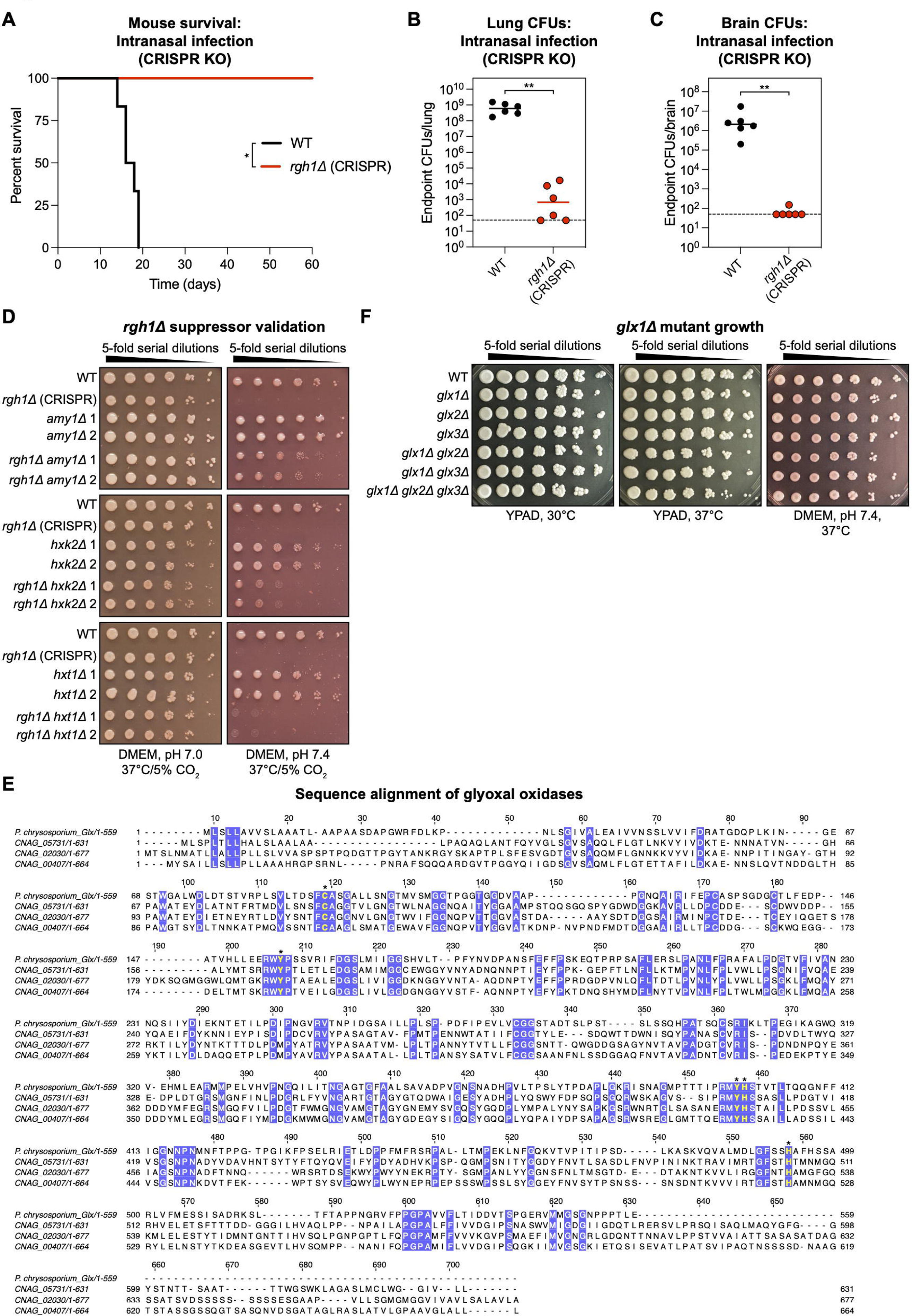
Additional data supporting roles for Rgh1 and glyoxal oxidases in fungal infectivity, related to Figures 6 and 7. (A) Survival of C57BL/6J mice infected intranasally with 5 x 10^4^ CFUs of either wild-type yeast or an independently constructed *rgh1Δ* mutant (*n* = 6 mice per group from 1 experiment). (B and C) Total endpoint CFUs isolated from the lungs (B) and brains (C) of mice in (A). Bars represent median values; dashed line represents limit of detection. Mice with undetectable CFUs were plotted at the limit of detection. (D) Spot dilution assays assessing genetic suppression of the *rgh1Δ* growth phenotype by candidates identified in Figure 6H. Images were taken after 2 days of growth. (E) Clustal Omega alignment of *C. neoformans* glyoxal oxidase protein sequences with the *Phanerochaete chrysosporium* glyoxal oxidase. Conserved catalytic residues are denoted in yellow with an asterisk. (F) Spot dilution assays assessing growth of glyoxal oxidase mutants. Images were taken after 3 days of growth. Statistical analyses were performed with Mantel-Cox (A) and Mann-Whitney (B, C) tests. \**p* < 0.05, \*\**p* < 0.01.

## Notes

### Competing Interest Statement

The authors have declared no competing interest.

### Summary of Updates

Abstract and Introduction have been updated

